# A multi-objective based clustering for inferring BCR clones from high-throughput B cell repertoire data

**DOI:** 10.1101/2021.10.01.462736

**Authors:** Nika Abdollahi, Anne de Septenville, Hugues Ripoche, Frédéric Davi, Juliana S. Bernardes

## Abstract

The adaptive B cell response is driven by the expansion, somatic hypermutation, and selection of B cell clones. A high number of clones in a B cell population indicates a highly diverse repertoire, while clonal size distribution and sequence diversity within clones can be related to antigen’s selective pressure. Identifying clones is fundamental to many repertoire studies, including repertoire comparisons, clonal tracking and statistical analysis. Several methods have been developed to group sequences from high-throughput B cell repertoire data. Current methods use clustering algorithms to group clonally-related sequences based on their similarities or distances. Such approaches create groups by optimizing a single objective that typically minimizes intra-clonal distances. However, optimizing several objective functions can be advantageous and boost the algorithm convergence rate. Here we propose a new method based on multi-objective clustering. Our approach requires V(D)J annotations to obtain the initial clones and iteratively applies two objective functions that optimize cohesion and separation within clones simultaneously. We show that under simulations with varied mutation rates, our method greatly improves clonal grouping as compared to other tools. When applied to experimental repertoires generated from high-throughput sequencing, its clustering results are comparable to the most performing tools. The method based on multi-objective clustering can accurately identify clone members, has fewer parameter settings and presents the lowest running time among existing tools. All these features constitute an attractive option for repertoire analysis, particularly in the clinical context to unravel the mechanisms involved in the development and evolution of B cell malignancies.

## 1 Introduction

Lymphocytes are the major cellular component of the adaptive immune system. They recognize and directly bind to pathogens (antigens) through a specific receptor, the B cell receptor (BCR), located on the outer cell surface. The BCR consists of two components: the recognition unit, structured by a membrane immunoglobulin (IG) protein, and an associated signaling unit. Upon antigen-specific stimulation, B lymphocytes undergo activation, proliferation, and further differentiation, enabling the secretion of the IG’s soluble form, known as antibodies. IG proteins are composed of two identical heavy chains (IGH) and two identical light chains (IGL). Each chain has two distinct parts: the variable region in the N-terminal side responsible for antigen recognition and the constant region on the C-terminal side anchored to the cell membrane. Complex genetic mechanisms create tremendous diversity of IGH and IGL variable regions. Three sets of genes encode variable regions in IGH: variable (V), diversity (D), and joining (J). These genes are naturally separated on the genome. However, they become juxtaposed during early B cell ontogeny by a process called VDJ recombination, which randomly selects and joins one of each of three types of genes together (*1*). Joining is imprecise as nucleotides are randomly deleted and inserted in the V-D (N1) and D-J (N2) junctions. The N1-D-N2 region is at the center of the so-called third complementarity determining region (in short, CDR3) and has the highest variability within the IGH molecule. VJ recombination also occurs for IGL genes, and random pairing of IGH and IGL chains allows the production of highly diverse B lymphocytes. In this work, we consider only IGH sequences since they are more diverse than IGL chains, providing a reliable signature for immune repertoire studies (*2*).

Upon antigen activation, B cells undergo rapid proliferation and further diversification of their BCR sequences by an enzymatically-driven process introducing nucleotide substitutions into the IG variable genes called somatic hypermutation (SHM). This occurs mainly in highly specialized structures, the germinal centers of secondary lymphoid organs, where a selection process operates termed affinity maturation. Cells for which SHM produced BCR with higher affinity for their cognate antigen expand, while those with a lower affinity are eliminated, thereby resulting in affinity maturation of the lymphocyte. Consequently, each B lymphocyte expresses a unique IG nucleotide sequence, enabling recognition of a particular set of antigens. The collection of an individual’s unique BCRs sequences is called BCR immune repertoire. During adaptive immune responses, B cell clones expand and contract. Such clones are important cellular units of the immune system. They consist of groups of B-cells derived from a common precursor during affinity maturation. Clonally-related B-cells share the same IGH and IGL rearrangements and can contain many molecular variants due to SHM.

Identifying clones in BCR repertoires (clonal grouping) is the starting point for several studies involving distinct subjects like autoimmune disease (*3*), cancer (*4*), and SARS-CoV-2 infection (*5*). Moreover, it is also a commonly used way to distinguish clonal (tumoral) from non-clonal (non-tumoral) B cell populations in case of suspicion of B cell malignancies (*6*). Extremely varied BCR repertoires are called polyclonal or non-clonal repertoires and are generally observed in healthy individuals. In contrast, individuals diagnosed with lymphoproliferative diseases such as B-cell leukemia or lymphoma have monoclonal or clonal BCR repertoires in which there is one highly expanded clone. In-between these two extreme situations, the immune repertoire can display unique or multiple minor clonal expansions reflecting various perturbations of the immune homeostasis such as infections, autoimmune diseases, and immunosenescence. This type of immune repertoire is termed oligoclonal.

Next-generation sequencing (NGS), which produces a large set of sequences, has profoundly changed our ability to study lymphocyte repertoires. Furthermore, identifying clones in high-throughput B cell repertoire data can help reconstruct cell lineage and unravel inter/intra clonal repertoire diversity. Several computational methods for clonal grouping have been developed, which generally employ clustering algorithms to infer clonal relationships (*7–9*). Most methods perform clonal grouping in two main steps. First, sequences with the same IGHV and IGHJ segments, and junctions of the same length, are grouped. Second, the sequences within each group are clustered according to some sequence-based distance. Any standard clustering approach can be applied, such as hierarchical (*10*), spectral (*8*) or agglomerative clustering (*7*). An alternative to these clustering approaches is to construct a lineage tree and cut it to create sub-trees, or clones (*11, 12*). All previous methods focus mainly on minimizing intra-clonal distances; they are based on only one criterion, reflecting a single measure of the partitioning quality. A single measure might not capture the different characteristics of the clustering problem, whereas a multi-objective approach might be more appropriate. Another possible limitation of existing methods is that such algorithms do not allow sequence interchanges once the groups are established.

Here we propose MobiLLe, a <u>M</u>ulti-<u>O</u>bjective <u>B</u>ased clustering for <u>I</u>nferring BCR clones from high-throughput B ce<u>LL</u> r<u>E</u>pertoire data. The method requires IGHV and IGHJ gene annotations and a fixed CDR3 identity threshold to form initial clones. Next, clones are refined by allowing sequences to move among different clusters until they find their appropriate place. For that, we optimize two objective functions for minimizing intra-clonal diversity and maximizing inter-clonal differences in parallel. Such functions continually evaluate clones’ consistency until no improvement is observed in their cohesion or separation. By minimizing intra-clonal diversity, we improve each clone’s cohesion, which measures how similar sequences are within the clone. On the other hand, by maximizing the inter-clonal differences, we improve the separation among distinct clones. We show that our approach greatly improves clonal grouping on simulated benchmarks and performs comparably to the most powerful and recent methods on experimental repertoires. MobiLLe produces reliable partitioning when existing clonal grouping inference methods fail, being very stable even on higher sequence mutation rates. When applied to experimental repertoires, When applied to experimental repertoires, it inferred clonal distributions similar to those of the most performing methods. However, MobiLLe has high scalability, low run-time, and minimal memory requirement. Moreover, it has a few parameter settings that do not require much effort and expertise to be tuned.

## 2 Methods

### 2.1 MobiLLe

Multi-objective clustering (MOC) decomposes a data set into related groups, maximizing multiple objectives in parallel. Several frameworks exist to implement MOC, MobiLLE relies on multi-run clustering, where a clustering algorithm runs multiple times to optimize different objectives that capture a compound fitness function (*13*). MobiLLE proceeds through two main steps: pre-clustering and refinement. Figure 1 shows its flowchart, and Algorithm 1 the pseudo-code for the refinement step.

**Figure 1:**
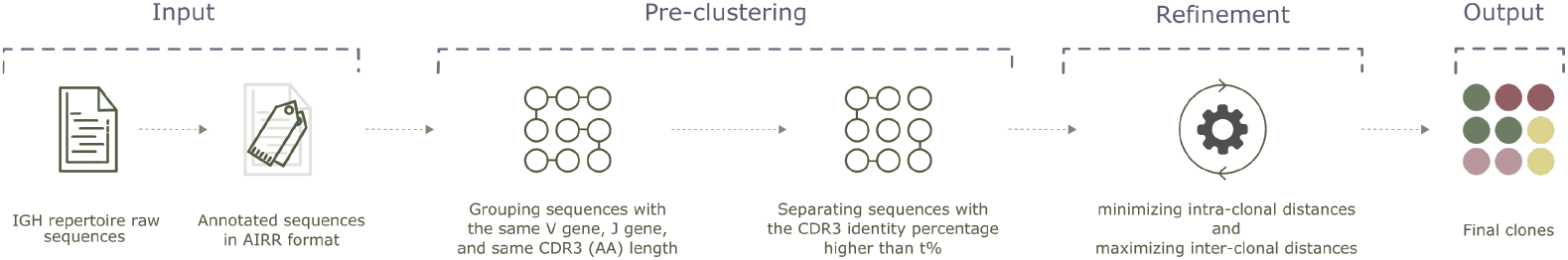
Flowchart of MobiLLe. The method requires IGH annotated sequences (IGHV, IGHJ, and CDR3 region were previously identified) to form initial clusters (pre-clustering step), we first group sequences with the same IGHV, IGHJ, and same CDR3 (AA) length; then, we separate sequences with less than t% CDR3 identity (default 70%). During the refinement step, sequences can move among different clusters until no improvement is observed in cluster cohesion or separation. The final groups represent clones with low intra-clonal diversity and high inter-clonal diversity.

#### 2.1.1 Pre-clustering

The pre-clustering step aims to group similar sequences to form initial clonal groups that can be refined later. First, sequences are annotated to identify their IGHV and IGHJ genes (and alleles) and locate their CDR3 regions. For this purpose, we used IMGT/HighV-QUEST (*14*), but in principle, any V(D)J annotation software could be used. Sequences with the same IGHV and IGHJ genes and the same CDR3 sequence length are then grouped together. Finally, we separate sequences with less than *t*% of CDR3 amino acid identity (by default *t* is 70%), see the “pre-clustering” panel in Figure 1.

#### 2.1.2 Clustering refinement

In this step, we iteratively refine clonal groups until we reach the minimum values for intra-clonal distances and the maximum values for inter-clonal distances. The algorithm described in 1 takes the initial clones *C* as input (generated during the pre-clustering step). For each sequence *i* ∈ *C* it computes two distances: *a_i_* (intra-clonal) and *b_i_* (inter-clonal). Such distances measure the cohesion/separation within detected clones; they were initially introduced to compute the Silhouette (*15*). *a_i_* is the average distance between the sequence *i* and any other sequence in the same clone; *b_i_* is the smallest average distance of *i* to all sequences in any other clone. In a well-detected cluster, *a_i_* is smaller than *b_i_*; thus, if for a given sequence *a_i_* is higher than *b_i_*, it might indicate that *i* was placed in a wrong cluster, and it should be moved to the cluster with the smallest average distance. If sequences are moved from a cluster *k* to a cluster *l*, then *a_i_* and *b_i_* need to be recomputed for all sequences in both clusters. Consequently, each sequence movement launches a new iteration of the algorithm, and it stops if no movement was observed in the previous iteration or after a predefined number of iterations.

Certainly, the distance metric *d*(*i,j*) (between sequences *i* and *j*) plays an important role when computing *a_i_* and *b_i_*. Distances based on sequence similarity of the whole sequences can be inaccurate since different IGHV, and IGHJ genes can present considerable similarities. Moreover, CDR3 regions are shorter than IGHV/IGHJ genes, and a normalized distance should be more appropriate. Therefore, we split the sequences into three parts, IGHV, IGHJ, and CDR3 region, and compute a different distance of each part, separately. The distance *d*(*i,j*) is the arithmetic mean of these three distances and is defined by the equation:

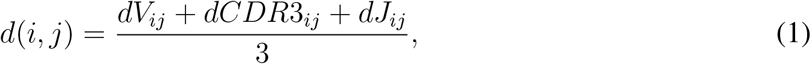

where *dV_ij_* is a binary distance based on IGHV gene identification, it is 0 if *i* and *j* were annotated with the same IGHV gene or 1 otherwise; dCDR3_*ij*_ is the normalized Levenshtein distance (*16*) between i’s and j’s CDR3 amino acid sequences; *dJ_ij_* is the normalized Levenshtein distance between i’s and j’s IGHJ nucleotide sequences. We recall that the Levenshtein distance computes the minimum number of single-character editions (insertions, deletions, or substitutions) required to transform one sequence into the other.

##### Algorithm 1: Clustering refinement

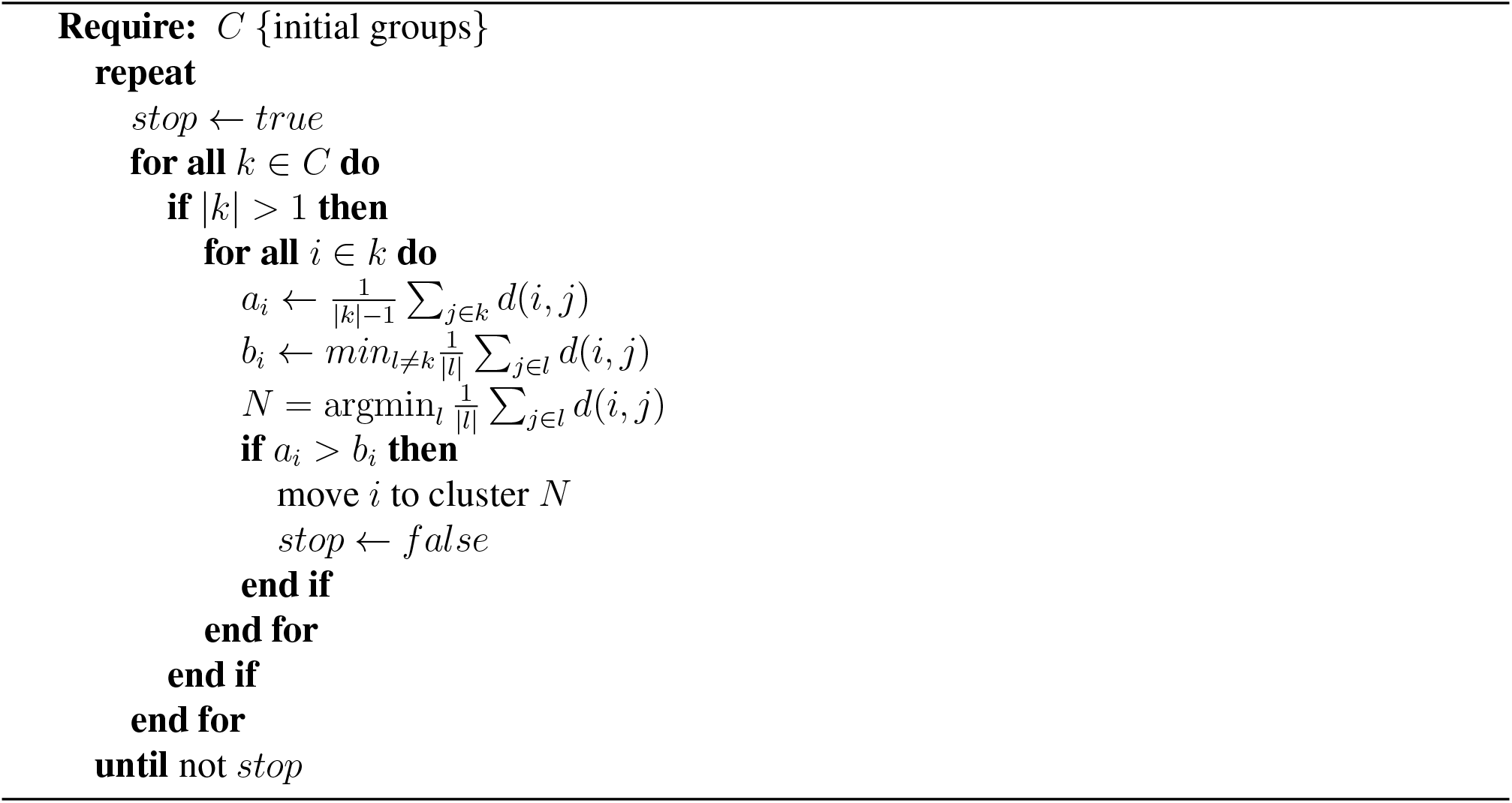

### 2.2 Data sets

To evaluate MobiLLe and compare it with existing clonal grouping methods, we used three types of BCR repertoire data: simulated, artificial, and experimental.

#### 2.2.1 Simulated repertoires

IGH simulated sequences are largely used to evaluate clonal grouping methods (*7, 8, 17*). Some repertoire simulators have been proposed as part of such tools, but, to the best of our knowledge, an independent B cell repertoire simulator that could produce different types of IGH repertoires (clonal and non-clonal) does not exist. In order to create simulated repertoires, we adapted GCtree (*18*), a B cell lineage simulator. We ran GCtree several times to produce independent B cell lineages that assembled represent a single repertoire.

To produce a B cell lineage, GCtree randomly selects IGHV, IGHD, and IGHJ germline genes from the IMGT database (*19*), then nucleotide(s) can be added to or removed from the IGHV-IGHD and IGHD-IGHJ junction regions. Next, a branching process is performed, and point mutations are included in the descendants. For the branching, GCtree uses an arbitrary offspring distribution that does not require an explicit bounding. Instead, it uses a Poisson distribution with parameter λ to estimate the expected number of offspring for each node. SHM are simulated by a sequence-dependent process, where mutations are preferentially introduced within certain hot-spot motifs. GCtree uses the 5-mer context model (*20*) to estimate the mutability *μ*_1_,…, *μ_i_*…, *μ_l_* for each residue *i* of a sequence of length *l*. The mutability of the whole sequence *μ*_0_, is then computed by averaging the mutability of its residues: 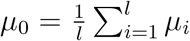. To determine the number of mutations *m* to be introduced in each mutant offspring sequence, GCtree also uses a Poisson distribution with parameter λ_0_, *m* is then computed as *Pois*(*μ*_0_λ_0_); note that more mutable sequences (higher *μ*_0_) tend to receive more point mutations.

Basically, GCtree simulator has two main parameters to be set: λ, to estimate the expected number of offspring of each node, and λ_0_, to determine the number of point mutations in mutant offspring sequences. We kept λ as the default value (e.g., 2), but we varied λ_0_ to produce simulations with different mutation rates. We experimented with four values {0.16, 0.26, 0.36, 0.46}, where 0.26 is the default value. Note that higher λ_0_ values produce more divergent B cell lineages. For each λ_0_ setting, we simulated three types of repertoires: monoclonal, oligoclonal, and polyclonal, obtaining 12 different simulations. The initial clonal size setting of each repertoire is shown in Table S1. Since we only kept productive sequences (without stop codons), the simulated repertoires can contain fewer sequences than expected.

#### 2.2.2 Artificial monoclonal repertoires

Gold standard experimental data, where truly clonal relationships are known with certainty, are difficult to obtain. In order to create experimental-based benchmarks, we constructed artificial monoclonal repertoires by combining sequences from the major clone of a monoclonal repertoire with sequences of a polyclonal repertoire. Our goal was to determine if clonal grouping methods could separate sequences from these two sources. To form a data set, we considered a total of 10000 sequences, where 10% of them were sampled from the major clone and 90% from the polyclonal background. Since we know the truly clonally-related sequences in each data set, we could compare the different tools for determining their grouping differences. We created three artificial repertoires from three different monoclonal samples, each having a specific V(D)J rearrangement. The major clone of the artificial monoclonal data set named AMR1 was annotated with IGHV1-69*01/IGHJ6*03 genes, the major clone of AMR2 with IGHV3-48*02/IGHJ4*02 rearrangements, and the AMR3 data set with the IGHV3-15*01/IGHJ6*02 rearrangements. We used IMGT/HighV-QUEST (*14*) as V(D)J assignment tool. AMR1, AMR2, and AMR3 major clone sampling were performed from monoclonal repertoires containing 22747, 20371, and 23665 sequences, respectively. The polyclonal background was sampled out of 136977 sequences.

Monoclonal samples and polyclonal background are human peripheral blood mononuclear cells obtained during routine diagnostic procedures at Pitié-Salpêtrière hospital (Paris-France). DNA sequences were obtained by polymerase chain amplification of IGH-VDJ rearrangements followed by NGS paired-end sequencing on an Illumina MiSeq platform. We obtained one “Read 1” and “Read 2” FASTQ files for each sample, which were then merged by the PEAR software (*21*). The merged FASTQ files were converted to FASTA format with seqtk (https://github.com/lh3/seqtk).

#### 2.2.3 Experimental repertoires

In order to evaluate our approach on realistically-sized data sets, we selected nine samples of human peripheral blood mono-nuclear cells collected during routine diagnostic procedures at Pitié-Salpêtrière hospital (Paris-France). From these repertoires, we peaked three samples to carry out a systematic comparison between MobiLLe’s results and existing clonal grouping methods. DNA sequences were obtained and assembled as described above. The first three columns in Table S2 show the number of reads (sequences), the number of unique sequences, and the clonality status defined by Genescan (*22*) (Figure S1) of each experimental repertoire.

### 2.3 Performance evaluation

#### 2.3.1 clustering accuracy

When clonal assignments were known, we could quantitatively assess clonal grouping algorithms’ ability in identifying clonally-related sequences. For that, we applied classical measures such as precision and recall for comparing the inferred clusters (clones) to the true ones. Consistently, we also computed the F-measure (FM), the harmonic mean of precision and recall; it is an aggregate measure of the inferred cluster’s quality. Precision and recall require three disjoint sets, which are: true positive (TP), false-positive (FP) and false-negative (FN). From these, we compute precision 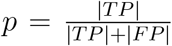, recall 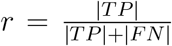, and FM 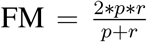. The values of these three metrics are in the interval [0, 1], being 1 the best and 0 the worst performance. Certainly, the way TP, FP and FN are computed will affect the accuracy measures. There are at least two ways to compute these values depending on the grouping level considered: *pairwise* and *closeness*.

The *pairwise* procedure considers the binary clustering task and focuses on the relationship between each pair of sequences. A pair of sequences is counted as: TP if the sequences are found together in both ‘true’ and ‘inferred’ clusters; FP if the sequences are found separated in the true, but together in the inferred clone; FN if the pair is found together in the true but separated in the inferred clone, see an illustration in Figure S2-A.

The *closeness* procedure evaluates clone compositions and the repertoire structure. For that, we first identified the best correspondence between inferred clones and correct clonal assignments. Then, we associated clone pairs that shared the maximum of common sequences. For each associated pair *i*, considering *I_i_* inferred and *T_i_* true clone, we computed TP_*i*_ as the intersection between the two sets (*I_i_* ∩ *T_i_*), FP_*i*_ as the difference between inferred and true clone (*I_i_* \ *T_i_*), and FN_*i*_ as the difference between true and inferred clone (*T_i_* \ *I_i_*). Finally, we computed 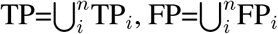 and 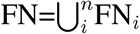, where *n* is the number of associated clone pairs; see an illustration in Figure S2-B.

#### 2.3.2 Comparison of clonal distributions

In order to compare clones obtained by different tools, we have defined five “events” that describe the differences between each pair of clonal distributions. For this, we labelled clusters in a clonal distribution *d*_2_ by comparing them with clusters in a distribution *d*_1_. These events are represented in Figure 2, and can be interpreted as follows:

1. identical: clusters in both distributions are identical, they contain the same set of sequences (Figure 2A),
2. join: when sequences of different clusters in *d*_1_ were joined in the same cluster in *d*_2_ (Figure 2B),
3. split: when sequences of a cluster in *d*_1_ were divided into several clusters in *d*_2_ (Figure 2C),
4. Mix: when a mixture of the join and split events occur. For instance, in Figure 2D, we observed two events, “split” (S_8_ and S_9_) and “join” (S_10_, S_11_ and S_12_),
5. Not found: when a cluster in *d*_2_ is not found among clusters in *d*_1_.

**Figure 2:**
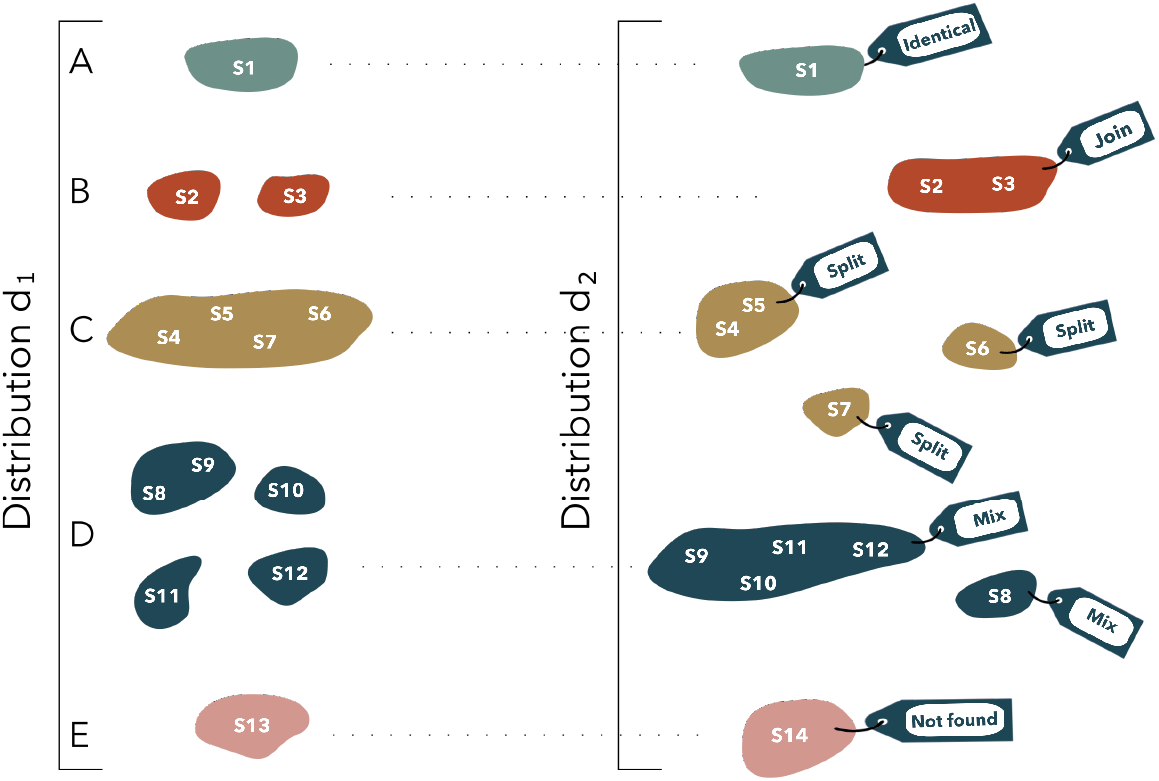
Clonal distribution comparisons. Five “events” describe the differences between two clonal distributions (*d*_1_ and *d*_2_) on the same set of sequences. The identical event counts the number of identical clones found in both distributions (a). The join event reports the number of clones in *d*_1_ found merged in *d*_2_ (b), while the split counts the number of clones in di found separated in *d*_2_ (c). The “mix” event is a mixture of these two later events (d) while “not found” reports the number of clones in *d*_2_ not found in *d*_1_ (e).

### 2.4 Clonal grouping tools considered for comparisons

MobiLLe groups, within a clone, sequences that might represent a B cell lineage. Thus, we chose four tools that employ the same clone definition. Each method has its own set of characteristics concerning the underlying algorithm, prior information, and produced outputs. Here we briefly describe them; for more details, report to original publications.

#### 2.4.1 BRILIA

B cell Repertoire Inductive Lineage and Immunosequence Annotator (BRILIA) builds up lineage tree reconstruction, clonal grouping, and V(D)J annotation into a single algorithm (*12*). From a collection of IGH sequences, BRILIA first provides initial V(D)J gene identification according to the IMGT database (*19*). Then, it groups sequences with the same IGHV and IGHJ gene subgroups and the same CDR3 sequence length. It proceeds by reconstructing lineage trees that will determine groups of clonally-related sequences. For that, it determines parent-child sequence relationships within each group for further reconstructing lineage trees. Evolutionary relationships are based on an adjusted hamming distance that penalizes dissimilarities in the N regions. Next, BRILIA determines the root of each obtained tree as the sequence involved in a cyclic dependency having the smallest distance to all other sequences in that cluster. Finally, a clone is a group of sequences sharing a common root sequence.

#### 2.4.2 Partis

Partis (*7*) uses hidden Markov models (HMMs) to represent V(D)J rearrangement events (*23*). An HMM is a probabilistic model, where the modelled system is assumed to be a Markov process with hidden states and unknown parameters. Each hidden state emits a symbol representing an elementary unit of the modelled data; for example, in BCR sequences, the hidden states represent either gene positions or N-region (addition or deletion) nucleotides. Thus, the HMM states represent nucleotides of each IGHV, IGHD, and IGHJ gene. The emission probabilities incorporate the probability of SHM at each nucleotide, and transition probabilities represent the probability of moving from one state to another. The HMM’ parameters (emission and transition probabilities) are estimated from a large panel of available sequences. Once the model is trained, BCR sequences are annotated by computing the Viterbi path through the HMM and finding the maximum-likelihood annotation. After V(D)J assignment, Partis applies its clonal grouping strategy. First, it creates initial clones of sequences sharing the same IGHV and IGHJ genes and the same CDR3 length. Then, it applies an agglomerative clustering algorithm to merge clusters that maximize the likelihood ratio that could indicate that two clusters derive from the same rearrangement events.

#### 2.4.3 SCOPe

SCOPe requires V(D)J annotation before clonal grouping, and tools such as IMGT/HighV-QUEST (*14*), or IgBlast (*24*) can be used. To identify a clone, SCOPe applies a spectral clustering method with an adaptive threshold to determine the local sequence neighbourhood; it means that it does not require a fixed threshold for detecting clonally-related sequences. Given a set of IGH sequences, SCOPe first divides them into groups with the same IGHV gene, IGHJ gene, and junction length. Then, it computes the similarity matrix for each group considering the hamming distance between junction regions of each pair of sequences within the group. Next, it generates a fully connected graph from the data points and performs local scaling to determine the local neighbourhood. Based on the graph, SCOPe builds an adjacency matrix and creates a Laplacian graph. The eigenvalues of such a graph can then be used to find the best number of clusters, and the eigenvectors can be used to find the actual cluster labels. Finally, SCOPe performs k-means clustering on the eigenvectors to get the labels (clone) for each node (sequence).

#### 2.4.4 SONAR

For SONAR (Software for the Ontogenic aNalysis of Antibody Repertoire) (*11*), a clonal group contains all IG reads that share a common ancestor. This tool focuses further on seeded lineage assignment, where the sequences of one or more known antibodies are used as seeds to find all sequences in the data set from the same lineage while leaving the rest of the sequences unclassified. In addition, it can perform “unseeded lineage assignment,” which consists of classifying sequences into component lineages without any additional information. In order to perform an unseeded lineage assignment, SONAR separates sequences based on their assigned IGHV and IGHJ genes. The sequences in each group are then clustered based on their CDR3 nucleotide identity (by default, 90% of CDR3 sequence), using the UCLUST algorithm in USEARCH (*25*). Eventually, each clone is identified as a distinct unseeded lineage.

## 3 Results

### 3.1 MobiLLe reconstructs simulated repertoires precisely

We first evaluated MobiLLe performance in reconstructing simulated repertoires with known clonal composition. The goal was to check if MobiLLe could accurately detect clonal memberships, clonal sizes, and distributions. In particular, we generated benchmarks that simulate several types of repertoire (clonal and non-clonal) with different SHM rates. For that, we adapted GCtree (*18*), a B cell lineage simulator, which randomly selects germline sequences for generating the unmutated common ancestor of each lineage and then introduces point mutations at hot-spot positions. Since GCtree generates a small number of productive sequences, we did not use selection models that could reduce the number of generated sequences. Nevertheless, we used a set of parameters to consider the following aspects of the B cell lineage biology: mutability (substitution), tree branching, and base-line mutation rates (Section 2.2.1).

We ran GCtree several times to create a collection of B cell lineages, composing a unique repertoire. To produce simulated repertoires with different SHM loads, we varied the corresponding GCtree parameter λ_0_ that determines the number of mutations in offspring sequences. We experimented with four different configurations {0.16, 0.26, 0.36, 0.46}, where higher values produce more divergent B cell lineages. We produced 12 simulated benchmarks, which simulate the three types of repertoires: monoclonal, oligoclonal, and polyclonal. The first four columns in Table 1 show SHM rates, clonality status, the number of sequences, and the number of expected clones for all simulated repertoires. Using these data, we evaluated our method by comparing inferred clones to truly related clonal sequences generated during the construction of each simulated repertoire. We used two complementary approaches in order to evaluate the clonal grouping accuracy: pairwise and closeness, detailed in Section 2.3.1.

**Table 1:**
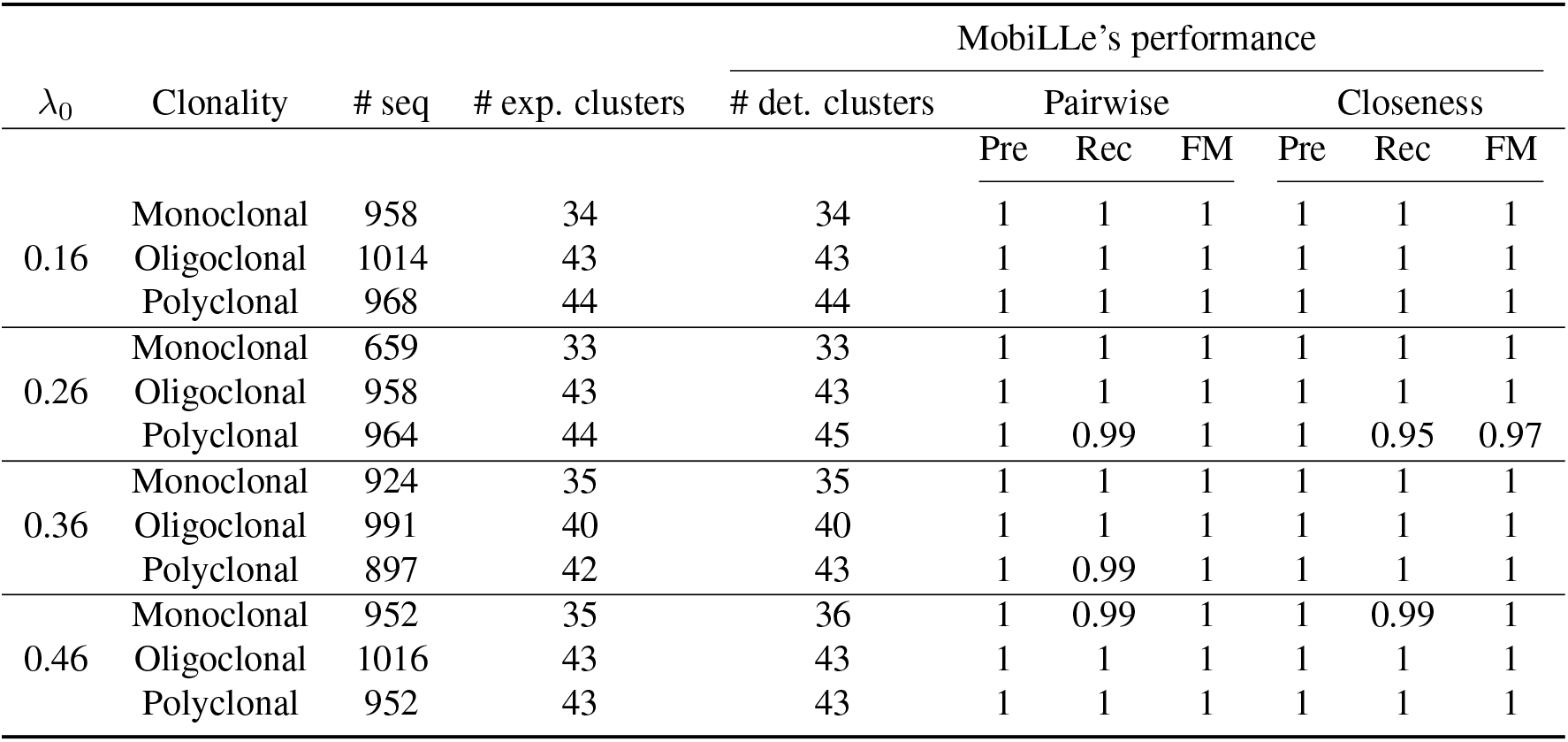
Evaluating the performance of MobiLLe on simulated repertoires. The third, fourth, and fifth columns show the number of sequences, the number of expected clones, and the number of detected clones, respectively. Pre, Rec, and FM are the abbreviations of precision, recall, and F-measure, respectively.

MobiLLe achieved high precision, recall, and F-measure across all simulated data sets for both pairwise and closeness performance measurement. Across all mutation rates, it accurately identified all pairwise relationships and reconstructed precisely the structure of all repertoires, see Table 1. The absolute performance measures were remarkably high for both clustering accuracy evaluation approaches, exhibiting a mean recall over 99%, and a mean precision/F-measure equal to 1. Closeness performance was lower (0.95) for polyclonal repertoire generated with λ_0_ = 0.26. For this data set MobiLLe inferred more clones than expected, increasing the number of false negatives and reducing the recall measure. MobiLLe detected the exact number of expected clones for the vast majority of repertoires, compare the fourth and fifth columns in Table 1.

### 3.2 Lower CDR3 identity thresholds do not impact the MobiLLe performance

In order to check the influence of MobiLLe parameter setting in the clonal grouping accuracy, we varied the pre-clustering threshold and measured the performance on simulated data sets. The MobiLLe’s pre-clustering step adopts three criteria to group clonally-related sequences: having the same IGHV gene/allele, the same IGHJ gene, and a CDR3 amino acid identity of at least t%. IGHV and IGHJ gene annotations are often used for grouping sequences into initial clusters by several tools (*7, 26*). The only controversial parameter is the CDR3 amino acid identity threshold. Many studies choose a fixed threshold and define any sequence with junction region sequence similarity above this threshold to be clonally-related (*3, 27*). We used a fixed threshold t=70% based on the definition of BCR subgroups with highly similar CDR3 motifs detailed in (*28*), often referred to as stereotyped BCR. However, other studies suggest different cutoffs, varying from 60% to 70% (*29–31*). To verify the impact of different cut-offs on the MobiLLe’s performance, we varied the pre-clustering threshold from 50% to 90%. The results are shown in Figure 3. For different repertoire types and different mutation rates, a threshold of 70% achieved the optimal performance. Note that a higher CDR3 identity threshold reduces the performance of MobiLLe, mainly in reconstructing the repertoire structure; see closeness performances (Figure 3BDF). It can be explained by the larger number of singletons generated with a higher threshold. Once singletons are formed, MobiLLe cannot merge them into higher density clusters since its intra-clonal distance *a_i_* is zero, and it is smaller than any other inter-clonal distance. We also observed that closeness performances degraded faster than pairwise when increasing CDR3 identity threshold (Figure 3BDF), mainly in the monoclonal repertoire with a high mutation rate (Figure 3B). This is due again to the effect of a large number of singletons that disturb the repertoire topology. On the other hand, a lower CDR3 identity threshold does not impact the MobiLLe performance significantly. Figures 3-ABCE show no influence, while Figures 3-DF present some performance perturbations. MobiLLe can adapt its performance when using CDR3 identity threshold in the 50%-70% range; higher values are not recommended since many singletons could be generated.

**Figure 3:**
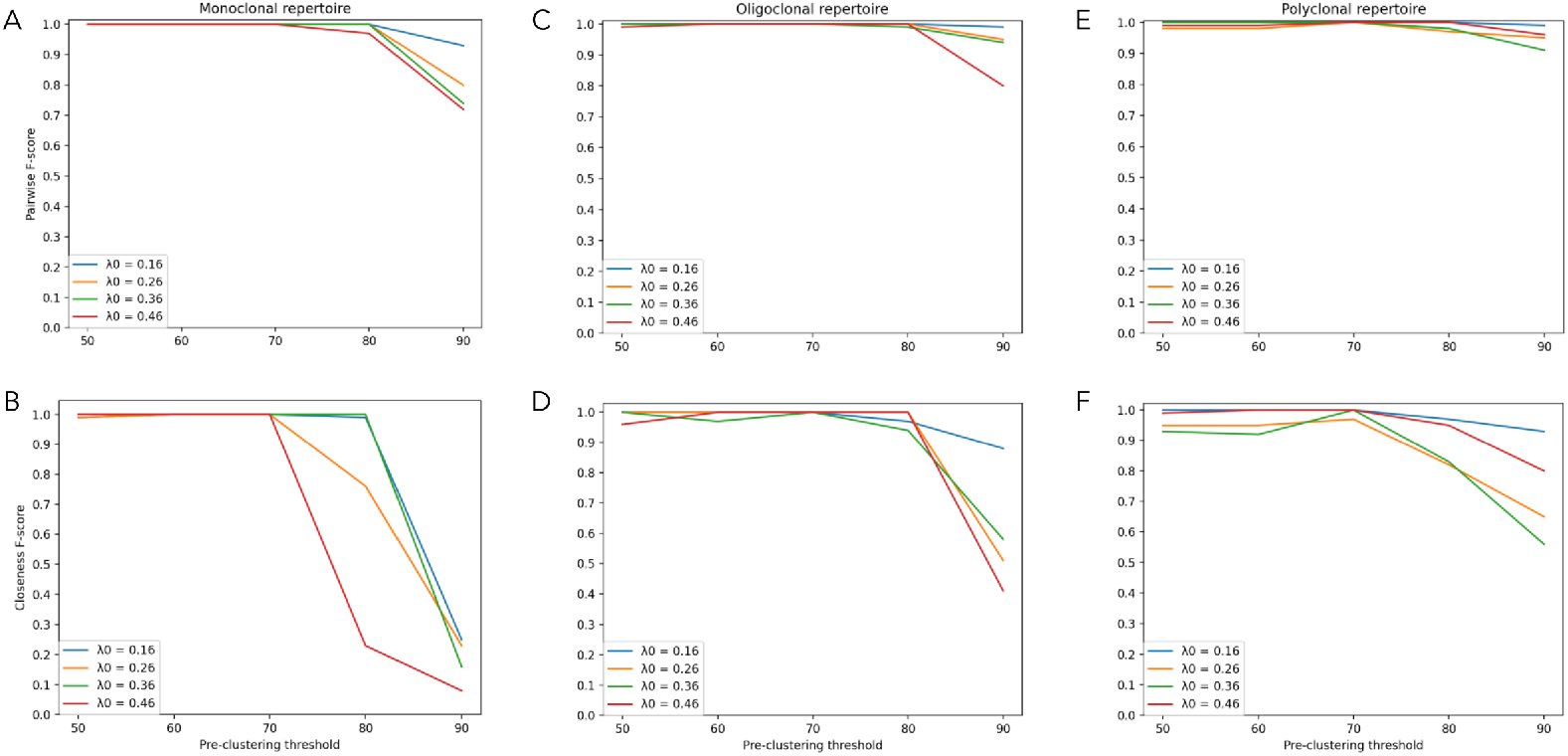
Effect of pre-clustering threshold on MobiLLe’s performance. We varied the pre-clustering threshold *t* from 50% to 90%, and evaluated the pairwise (top) and closeness (bottom) performance on 12 simulated repertoires with different SHM rates. For that we varied λ_0_ parameter of Gctree B cell lineage simulator.

### 3.3 Exploring the clonality of experimental repertoires

To demonstrate the application of MobiLLe on experimental data, we selected nine samples of human peripheral blood mononuclear cells collected during routine diagnostic procedures at the Pitié-Salpêtrière hospital. Three of these samples contained clonal leukemic cells, and six of them were considered non-clonal originating from patients devoid of malignancy (Section 2.2.3). Their clonality status was previously established by conventional methods, including PCR amplification of IGH-VDJ rearrangements followed by Genescan analysis (*22*) (Figure S1).

Figure 4 shows clonal abundances/densities for each analyzed sample. To measure the repertoire disequilibrium, we used the Gini index (*32*), which reflects the inequalities among values of a frequency distribution; zero indicates perfect equality, while one corresponds to maximal inequality. Clonal repertoires presented the highest Gini indexes, close to 1 for individuals 1 to 3 (see Figure 4ABC). Repertoires 1 and 3 presented similar clonal distributions, with the presence of a major clone accounting for the quasi-totality of the repertoire and a small number of minor clones having a low number of sequences (compare Figure 4A,C and Figures S3, and S5). MobiLLe results agreed with Genescan analysis, where we observed a single peak, indicating a monoclonal repertoire (Figure S1 A,C).

**Figure 4:**
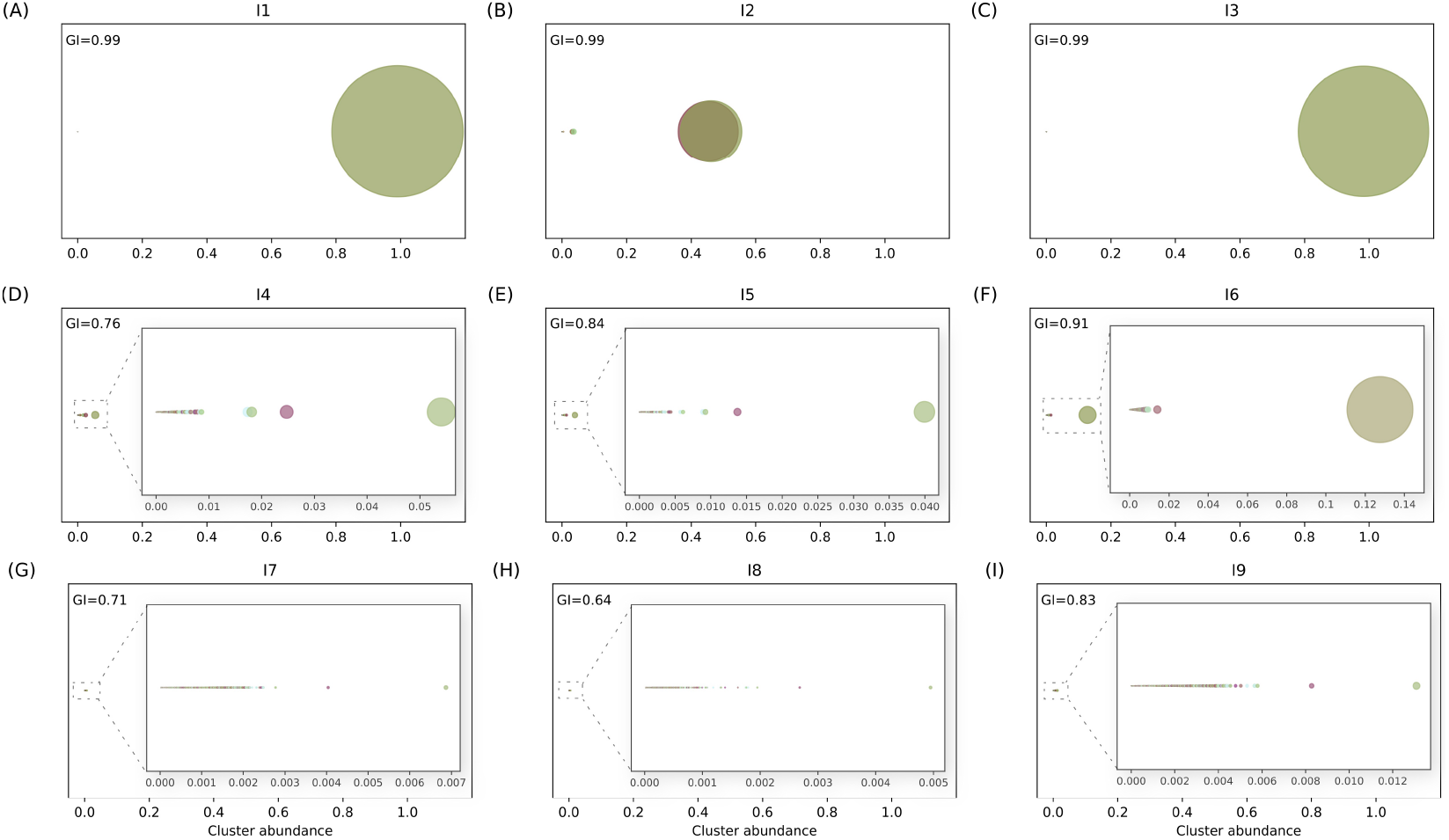
Clonal abundances/densities of nine experimental repertoires. Each circle represents a clone, and the clone’s abundance is displayed through its size. Gini index (GI) is an inequality index that measures clonal size distribution equitably; zero indicates perfect equality, while one corresponds to maximal inequality.

Individual 2 presented a different clonal profile with two major clones, each one accounting for more than 40% of the repertoire, see Figure 4B and Figure S4. Detailed sequence analysis revealed that the two major clones were composed of a productive and an unproductive IGH-VDJ rearrangement, corresponding to a leukemic cell population with biallelic IGH rearrangements. This biallelic IGH rearrangement was confirmed by Genescan analysis; Figure S1B shows two peaks, each corresponding to one IGH allele.

To better analyze non-clonal cases, we split them into two groups: (i) those with the predominance of clones with moderate abundances: *I*_4_, *I*_5_ and *I*_6_ (often termed oligoclonal repertoires), and (ii) the others with a more equilibrated (balanced) clonal distribution: *I*_7_, *I*_8_ and *I*_9_, termed as polyclonal repertoires. In the first group, repertoires 4 and 5 presented similar clonal profile, with Gini indexes around 0.76 and 0.84, respectively, see Figure 4DE and Figures S6 and S7. GeneScan analyses confirmed the non-clonal status of these individuals; we observed the absence of predominant peaks in Figures S1D,E. Individual 6 displayed a different configuration from other repertoires in the same group, with the presence of a relatively abundant clone representing 12% of the repertoire, Figure 4F and Figures S8. Not surprisingly, the repertoire 6 had the most biased distribution among non-clonal repertoires with a Gini index of 0.91. We also observed a more unbalanced profile in the Genescan analysis, Figure S1F.

In the second group, we found more homogeneous and less biased repertoires. Sample 7 and 8 had similar clonal distributions (Figure 4GH, Figures S9, and S10), while repertoire 9 was more irregular (Gini index = 0.83), see Figure 4I and Figure S11. We also observed a slight difference when comparing distributions generated by Genescan analysis, where repertoire 9 displayed more peaks than other repertoires in the same group, compare Figures S1-GHI. In these three cases, the size of the detected clones was relatively small, each of them accounting for less than 1% of the total sequences; see Figures S9, S10 and S11.

### 3.4 Comparison to existing clonal grouping methods

To obtain comparative results, we compared the performance of MobiLLe with four clonal grouping methods on both simulated and experimental data. We selected the most used tools: Partis, SCOPe, SONAR and BRILIA, briefly described in Section 2.4, and used the evaluation strategies described in Section 2.3 to measure their performances and compare clonal distributions.

#### 3.4.1 Simulated BCR repertoire data

We first compared MobiLLe to state-of-art methods on the 12 simulated repertoires described in Section 2.2.1. We used precision, recall and F-measure computed in two different ways, pairwise and closeness, to quantify clustering accuracy (Section 2.3.1).

##### Pairwise performances

MobiLLe achieved the best pairwise performance across all simulated data sets, Figure 5-Top. SCOPe and Partis outperformed SONAR and BRILIA. All tools achieved a precision close to 1, demonstrating that few false positives were detected. However, most tools had over-split clones, detecting many false negatives that considerably decreased recall and F-measure values.

**Figure 5:**
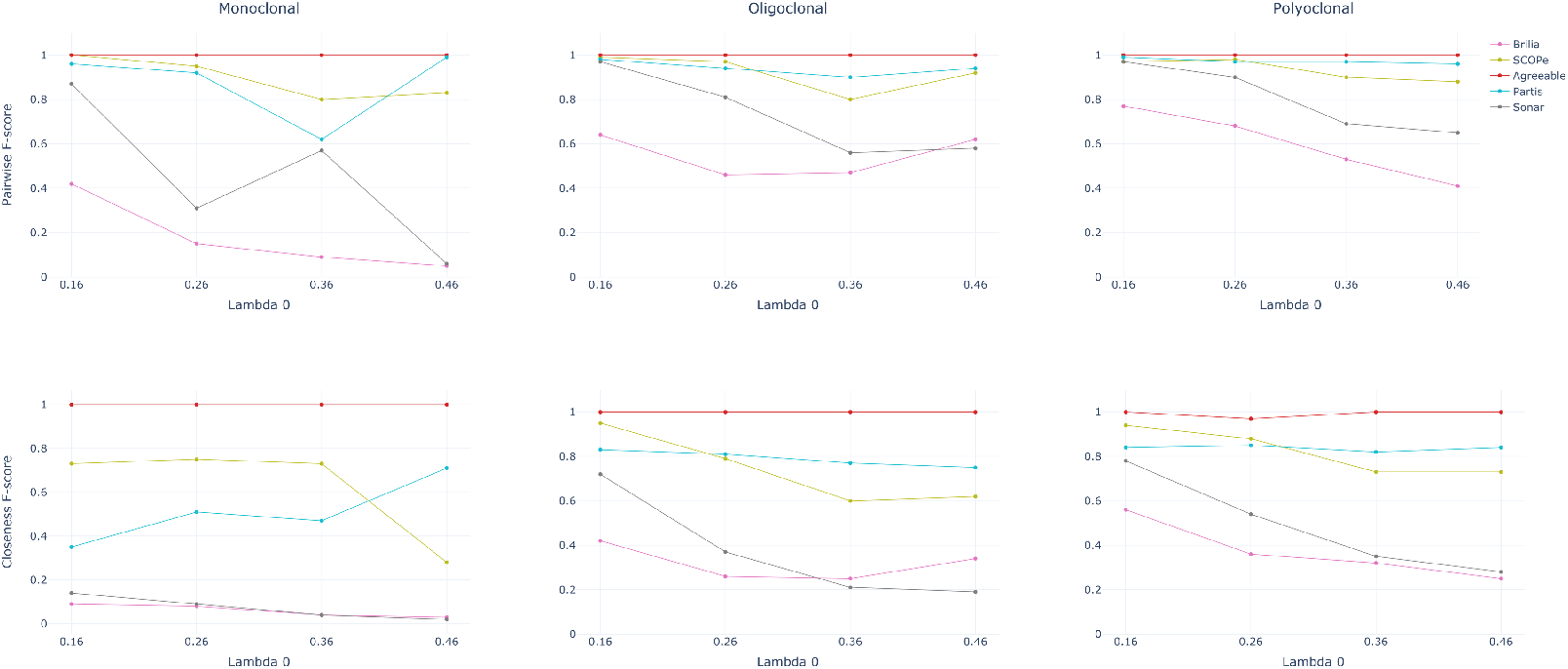
Comparison of clustering accuracy on simulated repertoires. We compared MobiLLe performances with four other tools: Partis, SCOPe, SONAR and BRILIA. For that, we generated benchmarks that simulated three types of repertoires (monoclonal, oligoclonal and polyclonal) with different SHM rates (0.16, 0.26, 0.36, 0.46), totalling 12 simulated samples. We used F-measure computed in two different ways, pairwise (top) and closeness (bottom), to quantify clustering accuracy.

SCOPe achieved high recall and F-measure for simulated data sets with lower mutation rates (λ_0_ = {0.16, 0.26}), see Tables S4-S9. Recall and F-measure values were above 0.94 for these six simulated repertoires. We observed lower recalls and F-measures for the remaining data sets, produced with higher mutation rates λ_0_ = {0.36, 0.46} (Tables S10-S15). On the other hand, Partis obtained a good pairwise performance across all simulated data sets independently of mutation rates. The only exception was the monoclonal repertoire produced with λ_0_ = 0.36 (Table S10). For this data set, Partis detected 44 clones when 35 were expected, decreasing its recall considerably. Interestingly, for lower mutation rates, SCOPe outperformed Partis, but we observed the opposite for higher mutation rates on most simulated repertoires. Thus, Partis seems to be more accurate when analyzing clonally-related sequences with higher divergence. Partis had difficulties in separating clusters with highly similar clonally-related sequences, as reported in the initial publication.

For oligoclonal and polyclonal repertoires, the different mutation rates seemed to influence SONAR performances. Recall and F-measure decreased as mutation rates increased, especially for the oligo-clonal repertoires. For the monoclonal samples with λ_0_ = 0.26 (Table S7), SONAR obtained lower recall and F-measure than with λ_0_ = 0.36 (Table S10). We observed that SONAR had over-split the largest clone of the first repertoire (λ_0_ = 0.26), grouping only 37% of sequences. On the other hand, it carried out less splits on the most abundant clone of monoclonal repertoire generated with λ_0_ = 0.36, grouping 62% of sequences. Once splits in large clones contribute to accuracy decreasing, it could explain the lower performance of SONAR on the monoclonal repertoire (λ_0_ = 0.26). For the monoclonal repertoire with λ_0_ = 0.46, SONAR detected four times more clones than expected, obtaining its lowest recall and F-measure, 0.03 and 0.06, respectively.

Most of the time BRILIA achieved the lowest pairwise performances across all simulated repertoires generated with different mutation rates. BRILIA removes sequences, which it cannot annotate, reducing the original data set, impacting the accuracy calculation. We also observed that BRILIA has over-split repertoires, producing the highest number of clones for most simulated data sets. The best performance was obtained on polyclonal repertoires generated with lower mutation rates (λ_0_ = {0.16, 0.26} - Tables S6-S9) and the lowest performance on monoclonal repertoires with higher mutation rates (λ_0_ = {0.36, 0.46} - Tables S10-S13).

##### Closeness performances

MobiLLe achieved the best closeness performance across all simulated data sets independently of mutation rates; see Figure 5-Bottom. The closeness evaluation approach tends to be more challenging than pairwise since clone properties such as size and distribution are also evaluated rather than pairwise relationships. Conversely, as observed for other tools, MobiLLe performance was not impacted by higher mutation rates or unbalanced clonal distributions. MobiLLe reconstructed all repertoire structure precisely, showing good stability and high accuracy. On the other hand, the four evaluated clonal grouping tools obtained high precision values but much lower recalls and F-measures. As observed for pairwise measures, SCOPe and Partis outperformed SONAR and BRILIA.

SCOPe outperformed Partis for most monoclonal repertoires, but Partis surpassed SCOPe for the majority of oligoclonal and polyclonal samples. SCOPe achieved higher F-measure values (> 0.73) on monoclonal repertoires generated with lower mutation rates (λ_0_ = {0.16, 0.26, 0.36} - Tables S4-S12). However, its performance sharply decreased on the monoclonal repertoire with the highest mutation rate, achieving 0.16 and 0.28 for recall and F-measure (Table S13). Its performances were affected by higher mutation rates; especially on oligoclonal and polyclonal samples, where we observed a notable difference between repertoires generated with λ_0_ = {0.16, 0.26}and those generated with λ_0_ = {0.36, 0.46}; Figure 5D,F. Higher mutation rates did not impact the performance of Partis on simulated repertoires. Its accuracy was stable on polyclonal repertoires and presented slight fluctuations on oligoclonal samples. Interesting, on the monoclonal data set, Partis achieved better performance for highly mutated repertoires; the best F-measure was obtained on the sample generated with λ_0_ = {0.46}; Figure 5B and Tables S4-S15.

SONAR performance was completely affected by higher mutation rates. We systematically observed lower recalls/F-measures as mutation rates increase (Figure 5DF and Tables S4-S15). Independently of mutation rates, SONAR achieved low performance on monoclonal repertoires; F-measures were smaller than 0.2 (Tables S4, S7, S10, S13). SONAR over-split the most abundant clones of monoclonal repertoires, which greatly decreased closeness performances. We observed better results on oligoclonal and polyclonal repertoires, especially on samples generated with lower mutation rates (λ_0_ = {0.16, 0.26} - Tables S4-S9). On the other hand, for all repertoires generated with higher mutation rates, SONAR achieved an F-measure inferior to 0.4 (Tables S10-S15). BRILIA achieved the lowest performance for most of the analyzed repertoires. The only exception was the oligoclonal sample generated with λ_0_ = 0.46, where it outperformed SONAR (Table S14). For the remaining data sets, SONAR overpassed BRILIA; we observed a notable difference mainly on repertoires generated with lower mutation rates, where BRILIA achieved low F-measure systematically (Tables S5, S6, S8, S9). BRILIA and SONAR achieved an equivalent performance in monoclonal repertoires, with very low values for recall and F-measure.

#### 3.4.2 Artificial experimental repertoires

To investigate the tools’ performance on experimental benchmarks, we created artificial experimental repertoires from BCR high throughput sequencing data as described in Section 2.2.2. We generated three artificial monoclonal repertoires (AMR1, AMR2, and AMR3) by mixing sequences from the major clone of a monoclonal repertoire (10%) and a polyclonal background (90%). Each data set contained 10k sequences, and the performance measured clonal grouping tools’ ability in identifying memberships within the most abundant clone. Accurate tools might group sequences from the major clone and separate in different clusters those from the polyclonal background. Thus, we counted the number of splits (SC) and false-positives (FP) in the major clone to measure the tools’ accuracy. We also used the alluvial diagram for visualizing clustering results; it represents flows between expected clones (left) and inferred ones (right).

Figure 6 shows the clonal grouping performance of different tools for AMR1, AMR2, and AMR3 data sets. Blue blocks represent the major clone (sampled from a monoclonal repertoire) in the alluvial diagrams and pink or orange inferred clones. Pink blocks contain only sequences belonging to the major clone (true positives), while the orange blocks contain sequences from the polyclonal background (false positives). Block height symbolizes the size of a clone, that is, the number of sequences. For AMR1, MobiLLe obtained the best clonal separation with no split (SC=0) and the smallest false positive (only 3). Partis and SCOPe also obtained fewer false positives but higher SC, 4 and 5, respectively. SONAR and BRILIA did not find any false positives, but both tools performed a significant number of splits, 90 and 64, respectively. Note that MobiLLe accurately reconstructed the whole lineage without detecting separations observed with other tools. The clonally-related sequences separated by other tools had different CDR3 lengths. They differ in the number of consecutive Tyrosine (Y) (ARDRRGEWPPSDYYYYYYMDV, ARDRRGEWPPSDYYYYMDV, and ARDRRGEWPPT-DYYYMDV). Since MobiLLe uses Levenshtein distance to compare CDR3 regions, it allows to group sequences with different lengths. In this case, MobiLLe tolerated the insertion/deletion of tyrosines because the rest of the sequences were similar and considered them as originating from the same lineage. Interestingly, during our NGS runs routine analyze tyrosines insertions/deletions have been observed frequently, mainly on BCR sequences annotated with IGHJ6 genes. The origin of such phenomenon (biological or sequencing artifact) remains unknown.

**Figure 6:**
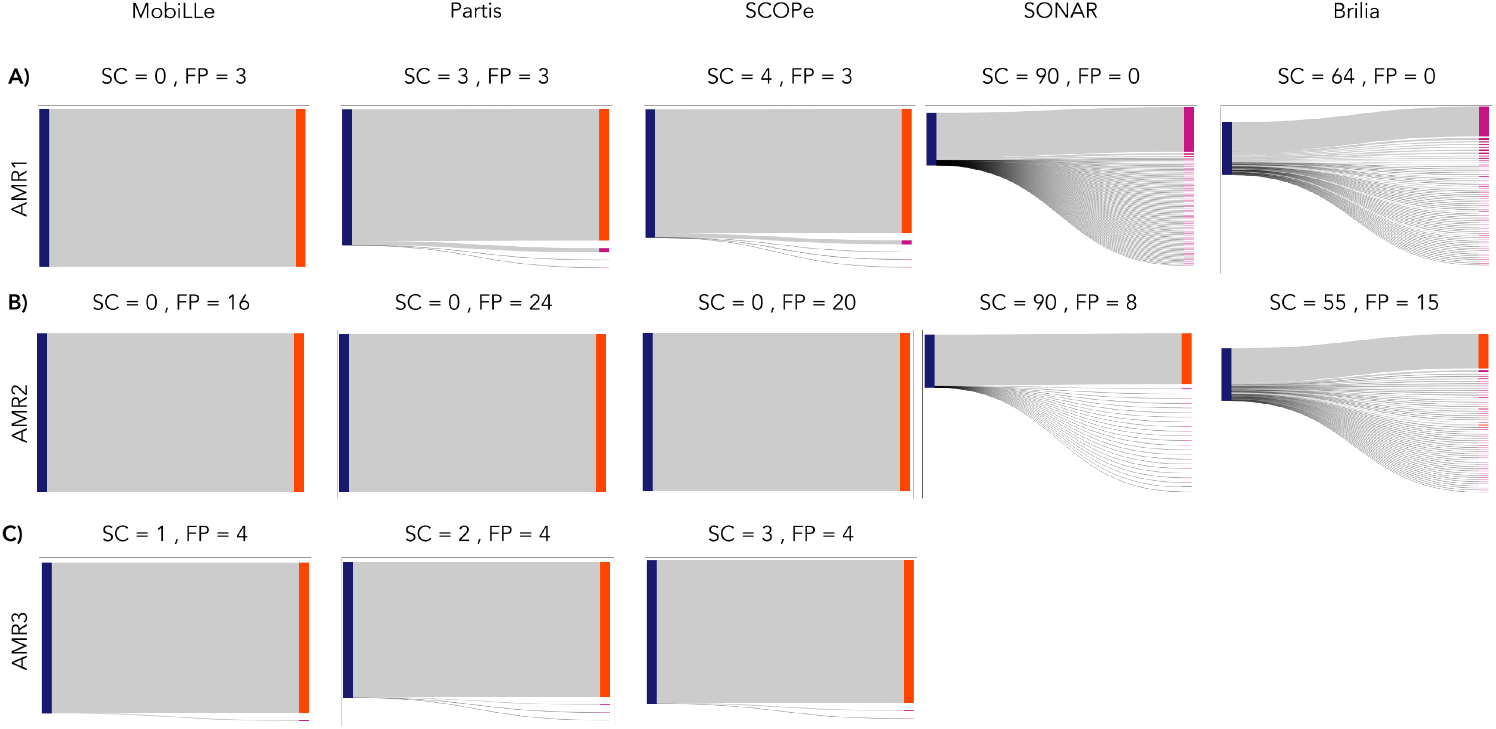
Performance comparison on artificial monoclonal repertoires. We generated three artificial monoclonal repertoires (AMR1, AMR2, and AMR3) by sampling sequences from the major clone of a monoclonal repertoire (10%) and a polyclonal background (90%). Each data set contained 10000 sequences. Accurate tools might group sequences from the major clone and separate those from the polyclonal background in different clusters. Therefore, we measured the performance of clonal grouping methods by computing the number of splits (SC) and false positives (FP) on the major clone. To better visualize and compare clustering results, we show alluvial diagrams for AMR1 (a), AMR2 (b) and AMR3 (c), where blue blocks represent the major clone and pink or orange inferred ones. Pink blocks contain only sequences belonging to the major clone (true positives), while the orange blocks contain sequences from the polyclonal background (false positives). SONAR and BRILIA did not produce results for the AMR3 data set since they do not deal with non-productive sequences.

As observed for AMR1, MobiLLe did not split the clonally-related sequences of AMR2 data set in different clones (SC = 0). It also obtained the lowest FP value. Similarly, Partis and SCOPe did not break the most abundant clone, but they detected more FPs than MobiLLe. SONAR and BRILIA still over-split the major clone, achieving an SC of 90 and 55, respectively. SONAR inferred more clones than BRILIA but obtained fewer FPs. Notably, the AMR3’s major clone contains non-productive sequences that SONAR and BRILIA did not consider. MobiLLe recovered almost all sequences of the most abundant clone (SC=1), followed by SCOPe (SC=2) and Partis (SC=3), see Figure 6C. The three tools detected the same number of FP=4. For all artificial monoclonal repertoires, we observed the same behavior in the results of each tool. We note that clonal grouping methods have clustered sequences differently. Partis, SCOPe, and MobiLLe have grouped the majority of sequences from the major clone of monoclonal repertoires. SONAR and BRILIA over-split clonal related sequences, but they detected fewer FPs. MobiLLe presented the best performance, achieving the minimum of splits and false positives.

#### 3.4.3 Clonal distribution comparisons

In order to understand the differences of clonal distributions of experimental repertoires, we compared MobiLLe clone composition to those of each considered clonal grouping tool. For comparing the methods, we selected three repertoires with different clonality status. I_1_ is a monoclonal repertoire with a major clone containing 98% of all sequences, I_2_ is a monoclonal repertoire with a biallelic IGH rearrangements, and I_8_ is a polyclonal repertoire; see Table S2 and Figure S1A,B,H.

We compared the inferred clones of existing clonal grouping tools with MobiLLe’s clustering results. For that, we used clustering quality measures and defined five events: identical, join, split, mix, and not found, which represent the (dis)similarities between two clonal distributions, see Section 2.3 and Figure 2. Table Table S16 and S17 show pairwise and closeness performance when comparing MobiLLe and other tools, while Figure 7 the occurrences of five events. Since MobiLLe removes sequences with no V(D)J annotations, we quantify the number of MobiLLe’s missing clones as “Not found” events. When analyzing the repertoire I_1_, we observed that BRILIA presented the maximum of identical clones, followed by Partis and SCOPe. These three tools obtained a similar number of clusters at the order of magnitude of the MobiLLe output. On the other hand, SONAR produced a significantly higher number of clusters, performing many splits. Interestingly, BRILIA and Partis achieved very high pairwise performances (Table S16), indicating that most pairs of clonally-related sequences were clustered in the same group. SCOPe inferred more clones than MobiLLe, achieving a slightly lower pairwise performance but around 0.98 (Table S16). Closeness comparison indicates that Partis clonal distribution is closer to MobiLLe than BRILIA and SCOPe (Table S17). However, the disagreement of these tools is mainly in the singleton identifications. On the other hand, pairwise and closeness recall/F-measure of SONAR was very low since it performed a considerable number of splits. All compared tools produced fewer joins and achieved high precision, meaning that they did not group sequences separated by MobiLLe.

**Figure 7:**
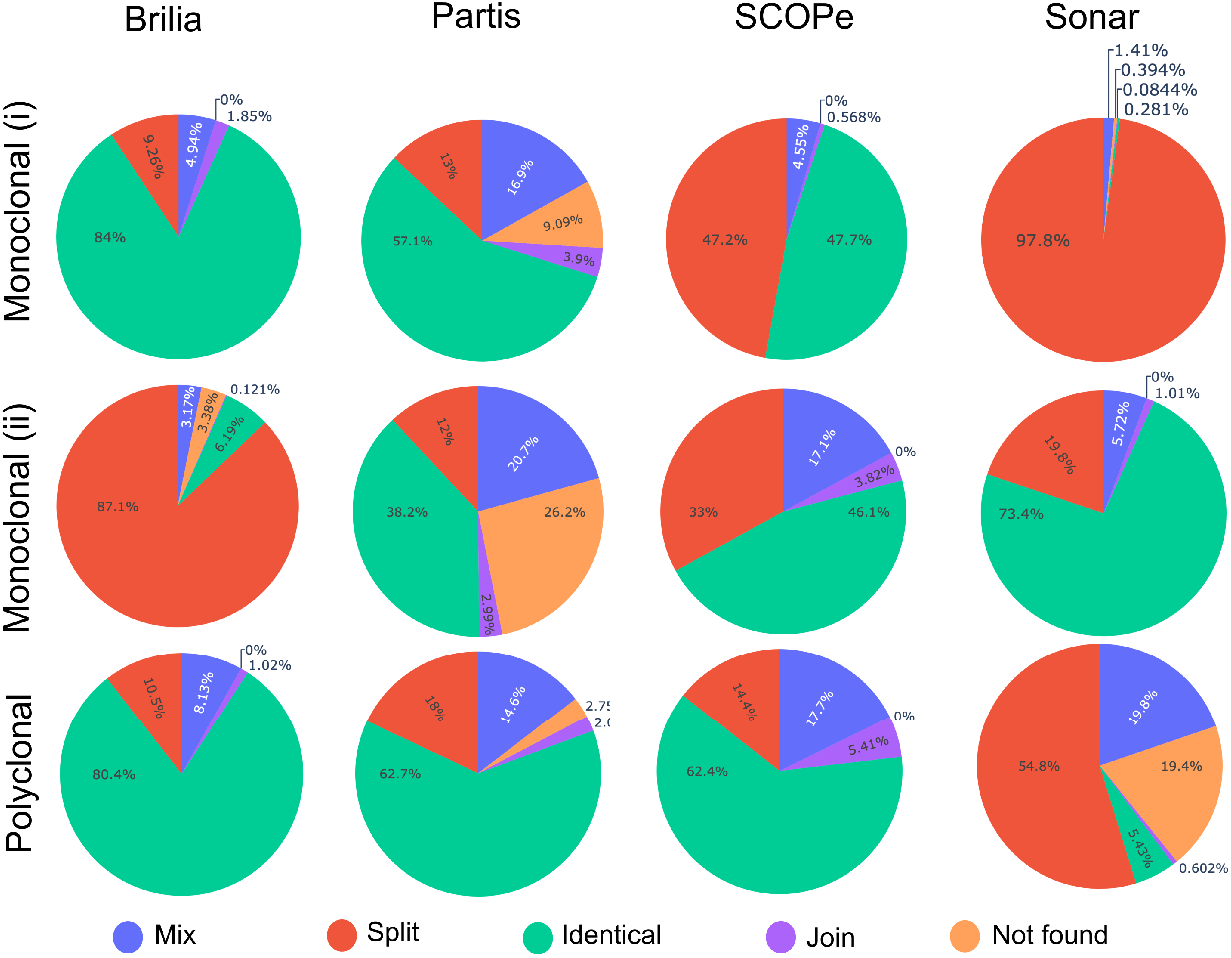
Clonal distribution comparisons on three experimental repertoires. We compared the inferred clones of each clonal grouping tool with MobiLLe’s clustering results. For that, we defined five events: identical, join, split, mix, and not found, representing the (dis)similarities between two clonal distributions: *d*_1_ (MobiLLe) and *d*_2_ (another tool). The “identical” event accounts for the percentage of identical clones found in both distributions; the “join” event reports the percentage of *d*_1_ clones found merged in *d*_2_ while “split” the percentage of *d*_1_ clones found separated in *d*_2_. The “mix” event accounts for a mixture of “join” and “split” events while “not found” reports the percentage of clones in *d*_2_ not found in *d*_1_; see an illustration in Figure 2.

For the biallelic monoclonal repertoire I_2_, SONAR inferred the highest number of identical clones, approximating the number of clones predicted by MobiLLe. For Partis and SCOPe, the predominant event was also “identical”, but both tools inferred fewer clones than MobiLLe. We also observed that the “mix” event was frequent, representing around 20%. Further analyses showed that “mix” event is associated with minor disagreement involving singletons. It was confirmed by pairwise performance values (Table S16) that show very high F-measures for SCOPe and Partis. SONAR achieved lower pairwise recall/F-measure than Partis and SCOPe, indicating more disagreements in the clonal distributions. Interestingly, SCOPe performed more splits than SONAR and Partis, but it did not impact its pairwise performances since most splits were not in the major clone as observed for SONAR. We also observed higher closeness values for Partis and SCOPe than SONAR. BRILIA over-split the data set I_2_, detecting three times more clones than MobiLLe. Consequently, its pairwise/closeness recall values were very low, only 0.11 (Table S16) and 0.01 (Table S17).

We observed more identical inferred clones in the polyclonal repertoire I_8_. BRILIA presented the highest value, accounting for more than 80%, followed by Partis and SCOPe around 62%. BRILIA and SCOPe achieved the best pairwise values. Although SCOPe inferred fewer clones than BRILIA, it identified most clonal membership correctly, achieving a comparable recall/F-measure. Both tools also presented the highest closeness F-measures. We observed a significant agreement between Partis and MobiLLe (pairwise and closeness values), but Partis performed more splits than BRILIA and SCOPe, which slightly decreased its pairwise/closeness recall. In general, the clonal composition of the three tools was similar to MobiLLe (Table S17). On the other hand, SONAR over-split the clones inferred by MobiLLe, detecting the largest number of groups. Moreover, its lower pairwise/closeness precision pinpointed many false positives predictions compared to MobiLLe results.

#### 3.4.4 Runtime

Computational efficiency is an important factor in clonal grouping development especially in a clinical context. Efficient methods need to process a large number of BCR sequences within a reasonable time without compromising clustering quality. In order to compare the computational requirements of MobiLLe with the four selected clonal grouping tools, we measured the running time on the I_1_, I_2_, and I_8_ repertoires. Figure 8 shows the time in seconds required by each tool to process the three repertoires, using a 3.4 GHz Octa-Core processor with 32 GB of memory; see the exact time in Table S18. MobiLLe took less than 20s to analyze the monoclonal repertoire I_1_, containing more than 30000 sequences. SONAR was also quick, but we observed an over-split that could explain its faster performance. BRILIA and SCOPe analyzed the I_1_ data set in a comparable time, around 200s and 100s, respectively. Partis was the most time-consuming tool, taking more than 2000s to process the I_1_ repertoire.

**Figure 8:**
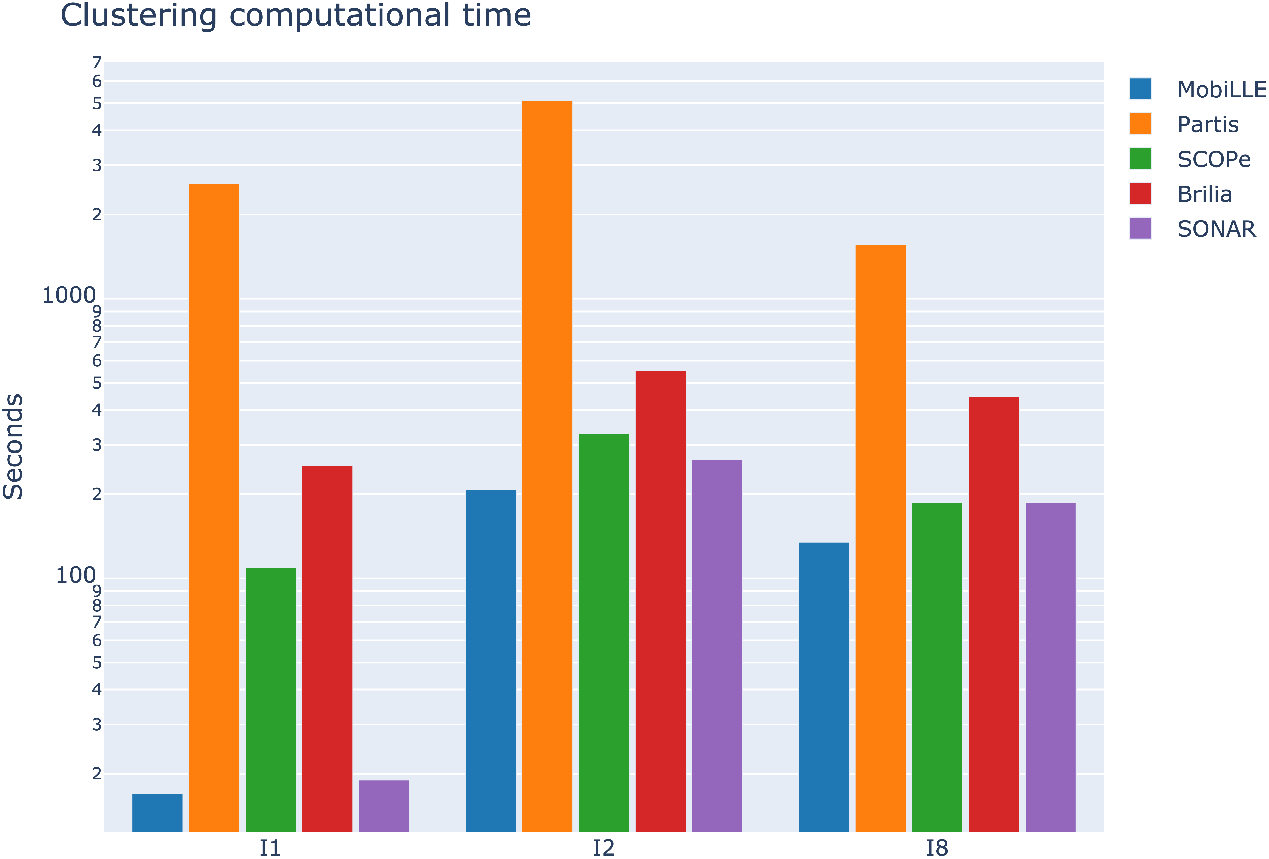
Comparing running times of clonal grouping tools. The running times for MobiLLe and other tools were measured for three experimental repertoires with different clonal compositions. To a better visualisation, we used log scale, Table S18 shows the time in seconds for each considered tool.

For I_2_, a biallelic monoclonal repertoire, containing more than 60000 sequences, MobiLLe also exhibited the fastest performance compared to the four evaluated tools. Of note, this sample took the longest time to be processed by all tools, including MobiLLe. Although samples I_1_ and I_2_ present a monoclonal structure, I_2_ has two major clones and significantly more clonal groups than I_1_. Apparently, the I_2_ structure slowed down the clustering processing since more comparisons among higher density clusters were done.

The polyclonal repertoire I_8_, containing more than 70000, seems less challenging than I_2_ with fewer sequences. Again, MobiLLe achieved the best time performance, clustering I_8_ in 135s. SONAR and SCOPe were also fast, taking 187s. Partis achieved the lowest performance, taking more than 1500s.

We observed that the clonal distribution significantly influences the running time of MobiLLe rather than the number of sequences. Calculating intra-clonal distances in repertoires with very abundant clones is more time-consuming. Accordingly, for repertoires with similar sizes but different clonal distributions, MobiLLe can present different running times. Therefore, for an equivalent number of sequences, the analysis of a monoclonal repertoire can be longer than the analysis of a polyclonal repertoire.

## 4 Discussion

The ability to obtain vast numbers (millions) of antigen receptor sequences using NGS techniques has dramatically changed our possibilities to explore immune (BCR) repertoires. Clonally-related sequences in a BCR repertoire descend from a common ancestor and present the same V(D)J rearrangement, but they may differ due to the accumulation of SHM, making their automatic clonal grouping challenging. Clonal relationships can be computationally identified from a large set of IGH sequences by using some clustering approach. Most clonal grouping methods automatically separate sequences into clonal groups based on their similarities or distances, considering the whole sequence and (or) junction regions. Commonly, their clustering algorithms are based on only one criterion that minimizes intra-clonal distances. However, such a single measure might not capture the different aspects of repertoires and considering multiple objectives seem more appropriate. Moreover, such algorithms do not review their clustering decisions, and sequences cannot move between clusters once the groups are established. Here we have proposed MobiLLe, a <u>M</u>ulti-<u>O</u>bjective <u>B</u>ased clustering for <u>I</u>nferring BCR clones from high-throughput B ce<u>LL</u> r<u>E</u>pertoire data. The method first produces initial clones containing sequences with identical IGHV and IGHJ germline annotations and more than t% of amino acid identity on CDR3 regions (default t=70%). Next, sequences can move among clones until achieving minimum intra-clonal diversity and maximum inter-clonal diversity. MobiLLe optimizes two objectives functions that continually evaluate clones’ consistency until no improvement is observed in their cohesion or separation.

We validated our method on synthetic data that simulated three types of immune repertoires (monoclonal, oligoclonal, and polyclonal) with different SHM rates. On the 12 simulated repertoires, MobiLLe inferred clonal relationships with very high accuracy, see Table 1, detecting clonal memberships and precisely reconstructed repertoire structures. The simulations showed that existing algorithms frequently over-split clones, separating sequences that belong to the same B cell lineage and originate from a common ancestor (Table S4-S15). High SHM rates impact the clustering accuracy of most methods; they achieved lower performance on simulated repertoires with divergent sequences, mainly on monoclonal samples. On the other hand, MobiLLe was very stable, keeping a good performance independently of SHM rates and repertoire types.

To evaluate performance on experimental data, we generated three artificial monoclonal repertoires by mixing sequences from the major clone of monoclonal repertoires with sequences from polyclonal repertoires. Then, we measured the ability of the existing clonal grouping tools and MobiLLe in grouping members of the most abundant clone. MobiLLe accurately grouped sequences from the major clone and separated in different clusters those from the polyclonal background. We observed that MobiLLe performed the minimum number of splits and detected fewer false positives compared with existing tools. In addition, our method can group sequences with different CDR3 lengths. This can be useful since SHM also introduces insertions and deletions (indels) in the junction region at a low-frequency (*33, 34*). It was the case of AMR1 data set where we observed insertion/deletion of tyrosine residues in the junction region. Thanks to this flexibility MobiLLe reconstructed the major clone of AMR1 perfectly.

After validating our method on simulated and experimental-based benchmarks, we applied it to experimental data sets where clonal groups were unknown. We selected nine repertoires, three of them contained clonal leukemic cells, and six were considered non-clonal (polyclonal). Clonality status was established a standardized methodology involving PCR amplification of IGH-VDJ rearrangements and Genescan analysis (*22*). We evaluated if MobiLLe could help estimate repertoires’ clonality and provide additional information about their clonal distribution. MobiLLe results agreed with Genescan analysis for all investigated samples (Figure S1), showing that it can predict clonality assignment and distinguish between clonal and non-clonal B cell populations. To help interpret clonal distributions, we have measured the disequilibrium of a repertoire with the Gini index applied to cluster size distribution. We observed that monoclonal repertoires presented the highest Gini indexes, indicating a disequilibrium in the clonal population. On the other hand, non-clonal repertoires exhibited lower Gini indexes, showing a more balanced data set. MobiLLe provides a collection of plots representing clones and their abundances for facilitating repertoire clonality interpretation. Compared to other clonal grouping tools, MobiLLe clonal distributions were often closer to Partis and SCOPe, the most performing tools. The pairwise evaluation confirmed that the clustering results of Partis and SCOPe were closer to MobiLLe with an F-measure superior to 0.93 for all analyzed repertoires. The clonal compositions of MobiLLe and BRILIA agreed on two samples but disagreed on the biallelic monoclonal repertoire. On the other hand, SONAR and MobiLLe agreed only on the biallelic monoclonal repertoire and disagreed on the other samples. MobiLLe was computationally more efficient than other tools. Moreover, it does not require an optimized distance threshold or training process, which can be time-consuming, especially when analyzing monoclonal repertoires.

Accurate identification of clonal members is essential for a wide range of repertoire analyses. However, the design and development of clonal grouping methods present several challenges, such as determining sequence similarities, choosing threshold distances, and maximizing computational efficiency. We show that MobiLLe clustering algorithm can identify clones with high accuracy and low runtime. We believe that our composed distance that analyses each sequence part is more appropriate and yields better results than previous definitions. A fixed distance threshold choice seems to be counterbalanced by our multi-objective optimization approach, which simultaneously optimizes the trade-off between intra-clonal cohesion and inter-clonal separability. Finally, MobiLLe is not as computationally demanding as methods that compute likelihood-based inference or optimized distance thresholds. The main weakness of our method is the impossibility of merging singletons to higher-density clusters. A possible improvement is to define a composed function that quantifies the clustering quality across all clones in the repertoire. Thereafter, merging singletons to other clusters might improve the objective function.

## 5 Conclusion

Clonal grouping is at the core of BCR repertoire analysis and all downstream investigations such as repertoire diversity estimation and intra-clonal analysis, among others, depending on the correct grouping of BCR sequences. Several clonal grouping methods have been proposed, but the most performing tools are either time-consuming or unstable when highly divergent repertoires are analysed. MobiLLe is a fast and accurate tool for clonal grouping with few parameter settings, low runtime, and memory usage that does not require a training process or hyper-parameter optimization. It can easily be applied to experimental repertoires of large sizes, proving useful plots to interpret BCR clonal groups.

## Author Contributions

NA, FD and JSB designed the project. NA developed the core algorithms and carried out the majority of the bioinformatic experiments. HR developed the code for SONAR analyses. AS and FD provided experimental data and analysed the results. NA and JSB wrote the manuscript. All authors reviewed, commented on, and approved the manuscript.

## Funding

This work has been supported by Sorbonne Universite grants: “2016 Programme Doctoral de Cancérologie” and SIRIC CURAMUS.

## Acknowledgments

Authors are grateful to Lucile Jeusset, Marine Armand and Thibaud Verny by insightful comments and fruitful discussion.

## Data Availability Statement

Data and the source code are freely available at github.com/julibinho/MobiLLe.

## Notes

https://github.com/julibinho/MobiLLe

### Competing Interest Statement

The authors have declared no competing interest.

