## Supplemental File for "A multi-objective based clustering for inferring BCR clones from high-throughput B cell repertoire data"

### A multi-objective based clustering for identifying clonally-related sequences from high-throughput B cell repertoire data
### Supplementary File

Nika Abdollahi, Anne Langlois De Septenville, Hugues Ripoche, Frédéric Davi and Juliana Bernardes

October 1, 2021

#### 1 Supplementary Tables

Table 1: **Initial clonal size distribution for the three types of simulated repertoires.** The clonal sizes were defined to guarantee that monoclonal repertoires will contain a major clone with at least 70% of sequences, oligoclonal repertoires will include two clones representing 14% and 9% of sequences% and any clone in polyclonal repertoires will not contain more than 5% of sequences.

| Monoclonal | | Oligoclonal | | Polyclonal | |
|---|---|---|---|---|---|
| #Clone | #sequences | #Clone | #sequences | #Clone | #sequences |
| 1 | 701 | 1 | 151 | 14 | 51 |
| 14 | 11 | 1 | 101 | 14 | 11 |
| 12 | 6 | 10 | 51 | 12 | 6 |
| 8 | 4 | 14 | 11 | 8 | 4 |
| 8 | 2 | 12 | 6 | 8 | 2 |
| | | 8 | 4 | | |
| | | 8 | 2 | | |
| 43 | 975 | 54 | 1036 | 56 | 988 |

Table 2: **Properties of experimental repertoires**. For each data set, we show the clonality status, a identifier, the total number of sequences and unique sequences, Gini index, number of total clones and those with at least two sequences

| Clonality | ind | # Sequences | # Unique Sequences | Gini coefficient | # clones | #clones wto singletons | % singletons |
|---|---|---|---|---|---|---|---|
| Monoclonal | 1 | 33599 | 22181 | 0.99 | 156 | 37 | 0.76 |
| | 2 | 65853 | 26431 | 0.99 | 304 | 69 | 0.77 |
| | 3 | 61990 | 19949 | 0.99 | 305 | 97 | 0.68 |
| Oligoclonal | 4 | 73888 | 53093 | 0.81 | 2343 | 1118 | 0.52 |
| | 5 | 294203 | 219006 | 0.85 | 19150 | 7801 | 0.59 |
| | 6 | 140026 | 84070 | 0.92 | 2398 | 654 | 0.72 |
| Polyclonal | 7 | 57076 | 40673 | 0.71 | 3493 | 1432 | 0.59 |
| | 8 | 70050 | 61379 | 0.64 | 11256 | 5160 | 0.54 |
| | 9 | 162742 | 104923 | 0.88 | 3955 | 1224 | 0.69 |

Table 3: **Evaluating the performance of MobiLLe on simulated repertoires**. The third, fourth, and fifth columns show the number of sequences, the number of expected clones, and the number of detected clones, respectively. Pre, Rec, and FM are the abbreviations of precision, recall, and F-measure, respectively.

| | | | | MobiLLe's performance | | | | | | |
|---|---|---|---|---|---|---|---|---|---|---|
| $\lambda_0$ | Clonality | # seq | # exp. clusters | # det. clusters | Pairwise | | | Closeness | | |
| | | | | | Pre | Rec | FM | Pre | Rec | FM |
| 0.16 | Monoclonal | 958 | 34 | 34 | 1 | 1 | 1 | 1 | 1 | 1 |
| | Oligoclonal | 1014 | 43 | 43 | 1 | 1 | 1 | 1 | 1 | 1 |
| | Polyclonal | 968 | 44 | 44 | 1 | 1 | 1 | 1 | 1 | 1 |
| 0.26 | Monoclonal | 659 | 33 | 33 | 1 | 1 | 1 | 1 | 1 | 1 |
| | Oligoclonal | 958 | 43 | 43 | 1 | 1 | 1 | 1 | 1 | 1 |
| | Polyclonal | 964 | 44 | 45 | 1 | 0.99 | 1 | 1 | 0.95 | 0.97 |
| 0.36 | Monoclonal | 924 | 35 | 35 | 1 | 1 | 1 | 1 | 1 | 1 |
| | Oligoclonal | 991 | 40 | 40 | 1 | 1 | 1 | 1 | 1 | 1 |
| | Polyclonal | 897 | 42 | 43 | 1 | 0.99 | 1 | 1 | 1 | 1 |
| 0.46 | Monoclonal | 952 | 35 | 36 | 1 | 0.99 | 1 | 1 | 0.99 | 1 |
| | Oligoclonal | 1016 | 43 | 43 | 1 | 1 | 1 | 1 | 1 | 1 |
| | Polyclonal | 952 | 43 | 43 | 1 | 1 | 1 | 1 | 1 | 1 |

Table 4: Monoclonal repertoire, generated with $\lambda_0 = 0.16$, number of sequences is equal to 958 and number of expected clusters is 34.

| | # analyzed seq | # clones | Pairwise | | | Closeness | | |
|---|---|---|---|---|---|---|---|---|
| Tool name | | | Precision | Recall | F-score | Precision | Recall | F-score |
| BRILIA | 922 | 68 | 1 | 0.26 | 0.42 | 1 | 0.05 | 0.09 |
| SONAR | 958 | 52 | 1 | 0.77 | 0.87 | 1 | 0.07 | 0.14 |
| Partis | 958 | 43 | 1 | 0.92 | 0.96 | 1 | 0.21 | 0.35 |
| SCOPe | 958 | 35 | 1 | 0.99 | 1 | 1 | 0.58 | 0.73 |
| MobiLLe | 958 | 34 | 1 | 1 | 1 | 1 | 1 | 1 |

Table 5: Oligoclonal repertoire, generated with $\lambda_0 = 0.16$, number of sequences is equal to 1014 and number of expected clusters is 43.

| | # analyzed seq | # clones | Pairwise | | | Closeness | | |
|---|---|---|---|---|---|---|---|---|
| Tool name | | | Precision | Recall | F-score | Precision | Recall | F-score |
| BRILIA | 658 | 69 | 1 | 0.47 | 0.64 | 1 | 0.26 | 0.42 |
| SONAR | 1014 | 55 | 1 | 0.94 | 0.97 | 1 | 0.56 | 0.72 |
| Partis | 1014 | 52 | 1 | 0.96 | 0.98 | 1 | 0.71 | 0.83 |
| SCOPe | 1014 | 46 | 1 | 0.97 | 0.99 | 1 | 0.9 | 0.95 |
| MobiLLe | 1014 | 43 | 1 | 1 | 1 | 1 | 1 | 1 |

Table 6: Polyclonal repertoire, generated with $\lambda_0 = 0.16$, number of sequences is equal to 968 and number of expected clusters is 44.

| | # analyzed seq | # clones | Pairwise | | | Closeness | | |
|---|---|---|---|---|---|---|---|---|
| Tool name | | | Precision | Recall | F-score | Precision | Recall | F-score |
| BRILIA | 947 | 78 | 1 | 0.63 | 0.77 | 1 | 0.39 | 0.56 |
| SONAR | 968 | 55 | 1 | 0.93 | 0.97 | 1 | 0.64 | 0.78 |
| Partis | 968 | 53 | 1 | 0.97 | 0.99 | 1 | 0.72 | 0.84 |
| SCOPe | 968 | 46 | 1 | 0.94 | 0.97 | 1 | 0.88 | 0.94 |
| MobiLLe | 968 | 44 | 1 | 1 | 1 | 1 | 1 | 1 |

Table 7: Monoclonal repertoire, generated with $\lambda_0 = 0.26$, number of sequences is equal to 659 and number of expected clusters is 33.

| Tool name | # analyzed seq | # clones | Pairwise Precision | Pairwise Recall | Pairwise F-score | Closeness Precision | Closeness Recall | Closeness F-score |
|---|---|---|---|---|---|---|---|---|
| BRILIA | 618 | 72 | 1 | 0.08 | 0.15 | 1 | 0.04 | 0.08 |
| SONAR | 659 | 72 | 1 | 18 | 0.31 | 1 | 0.05 | 0.09 |
| Partis | 659 | 42 | 1 | 0.85 | 0.92 | 1 | 0.34 | 0.51 |
| SCOPe | 659 | 36 | 1 | 0.9 | 0.95 | 1 | 0.6 | 0.75 |
| MobiLLe | 659 | 33 | 1 | 1 | 1 | 1 | 1 | 1 |

Table 8: Oligoclonal repertoire, generated with $\lambda_0 = 0.26$, number of sequences is equal to 958 and number of expected clusters is 43.

| Tool name | # analyzed seq | # clones | Pairwise Precision | Pairwise Recall | Pairwise F-score | Closeness Precision | Closeness Recall | Closeness F-score |
|---|---|---|---|---|---|---|---|---|
| BRILIA | 745 | 90 | 1 | 0.29 | 0.46 | 1 | 0.15 | 0.26 |
| SONAR | 958 | 87 | 1 | 0.67 | 0.81 | 1 | 0.22 | 0.37 |
| Partis | 958 | 52 | 1 | 0.88 | 0.94 | 1 | 0.69 | 0.81 |
| SCOPe | 958 | 50 | 1 | 0.94 | 0.97 | 1 | 0.65 | 0.79 |
| MobiLLe | 958 | 43 | 1 | 1 | 1 | 1 | 1 | 1 |

Table 9: Polyclonal repertoire, generated with $\lambda_0 = 0.26$, number of sequences is equal to 964 and number of expected clusters is 44.

| Tool name | # analyzed seq | # clones | Pairwise Precision | Pairwise Recall | Pairwise F-score | Closeness Precision | Closeness Recall | Closeness F-score |
|---|---|---|---|---|---|---|---|---|
| BRILIA | 876 | 109 | 1 | 0.51 | 0.68 | 1 | 0.22 | 0.36 |
| SONAR | 964 | 79 | 1 | 0.82 | 0.9 | 1 | 0.37 | 0.54 |
| Partis | 964 | 53 | 1 | 0.94 | 0.97 | 1 | 0.74 | 0.85 |
| SCOPe | 964 | 49 | 1 | 0.95 | 0.98 | 1 | 0.78 | 0.88 |
| MobiLLe | 964 | 45 | 1 | 0.99 | 1 | 1 | 0.95 | 0.97 |

Table 10: Monoclonal repertoire, generated with $\lambda_0 = 0.36$, number of sequences is equal to 924 and number of expected clusters is 35.

| Tool name | # analyzed seq | # clones | Pairwise Precision | Pairwise Recall | Pairwise F-score | Closeness Precision | Closeness Recall | Closeness F-score |
|---|---|---|---|---|---|---|---|---|
| BRILIA | 897 | 116 | 1 | 0.04 | 0.09 | 1 | 0.02 | 0.04 |
| SONAR | 924 | 105 | 1 | 0.40 | 0.57 | 1 | 0.02 | 0.04 |
| Partis | 924 | 44 | 1 | 0.45 | 0.62 | 1 | 0.31 | 0.47 |
| SCOPe | 924 | 36 | 1 | 0.66 | 0.8 | 1 | 0.58 | 0.73 |
| MobiLLe | 924 | 35 | 1 | 1 | 1 | 1 | 1 | 1 |

Table 11: Oligoclonal repertoire, generated with $\lambda_0 = 0.36$, number of sequences is equal to 991 and number of expected clusters is 40.

| Tool name | # analyzed seq | # clones | Pairwise Precision | Pairwise Recall | Pairwise F-score | Closeness Precision | Closeness Recall | Closeness F-score |
|---|---|---|---|---|---|---|---|---|
| BRILIA | 724 | 102 | 1 | 0.3 | 0.47 | 1 | 0.14 | 0.25 |
| SONAR | 991 | 124 | 1 | 0.38 | 0.56 | 1 | 0.12 | 0.21 |
| Partis | 991 | 49 | 1 | 0.81 | 0.9 | 1 | 0.62 | 0.77 |
| SCOPe | 991 | 57 | 0.99 | 0.66 | 0.8 | 1 | 0.43 | 0.60 |
| MobiLLe | 991 | 40 | 1 | 1 | 1 | 1 | 1 | 1 |

Table 12: Polyclonal repertoire, generated with $\lambda_0 = 0.36$, number of sequences is equal to 897 and number of expected clusters is 42.

| | # analyzed seq | # clones | Pairwise | | | Closeness | | |
|---|---|---|---|---|---|---|---|---|
| Tool name | | | Precision | Recall | F-score | Precision | Recall | F-score |
| BRILIA | 639 | 101 | 1 | 0.35 | 0.53 | 1 | 0.19 | 0.32 |
| SONAR | 897 | 120 | 1 | 0.53 | 0.69 | 1 | 0.21 | 0.35 |
| Partis | 897 | 51 | 1 | 0.94 | 0.97 | 1 | 0.7 | 0.82 |
| SCOPe | 897 | 58 | 1 | 0.82 | 0.9 | 1 | 0.57 | 0.73 |
| MobiLLe | 897 | 43 | 1 | 0.99 | 1 | 1 | 1 | 1 |

Table 13: Monoclonal repertoire, generated with $\lambda_0 = 0.46$, number of sequences is equal to 952 and number of expected clusters is 35.

| | # analyzed seq | # clones | Pairwise | | | Closeness | | |
|---|---|---|---|---|---|---|---|---|
| Tool name | | | Precision | Recall | F-score | Precision | Recall | F-score |
| BRILIA | 926 | 152 | 1 | 0.02 | 0.05 | 1 | 0.01 | 0.03 |
| SONAR | 952 | 157 | 1 | 0.03 | 0.06 | 1 | 0.01 | 0.02 |
| Partis | 952 | 44 | 1 | 0.97 | 0.99 | 1 | 0.55 | 0.71 |
| SCOPe | 952 | 46 | 1 | 0.73 | 0.83 | 1 | 0.16 | 0.28 |
| MobiLLe | 952 | 36 | 1 | 0.99 | 1 | 1 | 0.99 | 1 |

Table 14: Oligoclonal repertoire, generated with $\lambda_0 = 0.46$, number of sequences is equal to 1016 and number of expected clusters is 43.

| | # analyzed seq | # clones | Pairwise | | | Closeness | | |
|---|---|---|---|---|---|---|---|---|
| Tool name | | | Precision | Recall | F-score | Precision | Recall | F-score |
| BRILIA | 689 | 96 | 1 | 0.44 | 0.62 | 1 | 0.2 | 0.34 |
| SONAR | 1016 | 160 | 1 | 0.40 | 0.58 | 1 | 0.1 | 0.19 |
| Partis | 1016 | 53 | 1 | 0.89 | 0.94 | 1 | 0.6 | 0.75 |
| SCOPe | 1016 | 62 | 1 | 0.84 | 0.92 | 1 | 0.45 | 0.62 |
| MobiLLe | 1016 | 43 | 1 | 1 | 1 | 1 | 1 | 1 |

Table 15: Polyclonal repertoire, generated with $\lambda_0 = 0.46$, number of sequences is equal to 952 and number of expected clusters is 43.

| | # analyzed seq | # clones | Pairwise | | | Closeness | | |
|---|---|---|---|---|---|---|---|---|
| Tool name | | | Precision | Recall | F-score | Precision | Recall | F-score |
| BRILIA | 705 | 123 | 1 | 0.26 | 0.41 | 1 | 0.14 | 0.25 |
| SONAR | 952 | 147 | 1 | 0.48 | 0.65 | 1 | 0.17 | 0.28 |
| Partis | 952 | 52 | 1 | 0.92 | 0.96 | 1 | 0.72 | 0.84 |
| SCOPe | 952 | 57 | 1 | 0.79 | 0.88 | 1 | 0.57 | 0.73 |
| MobiLLe | 952 | 43 | 1 | 1 | 1 | 1 | 1 | 1 |

Table 16: Comparison of MobiLLE with four different clonal grouping methods on the three experimental data sets, using **pairwise evaluation method.**

| | # seq | # clones by MobiLLe | | # analyzed seq | # clones | Precision | Recall | F-score |
|---|---|---|---|---|---|---|---|---|
| $I_1$ | 33599 | 162 | BRILIA | 33578 | 161 | 0.99 | 0.99 | 0.99 |
| | | | SONAR | 29335 | 3542 | 0.99 | 0.21 | 0.36 |
| | | | Partis | 33585 | 73 | 0.99 | 0.99 | 0.99 |
| | | | SCOPe | 33554 | 176 | 0.99 | 0.96 | 0.98 |
| $I_2$ | 140026 | 2398 | BRILIA | 90236 | 7234 | 0.99 | 0.11 | 0.2 |
| | | | SONAR | 138439 | 1975 | 0.99 | 0.87 | 0.93 |
| | | | Partis | 138688 | 927 | 0.99 | 0.99 | 1 |
| | | | SCOPe | 138250 | 1387 | 0.99 | 0.98 | 0.99 |
| $I_8$ | 70050 | 11256 | BRILIA | 68133 | 10461 | 0.96 | 0.90 | 0.93 |
| | | | SONAR | 68171 | 17096 | 0.57 | 0.04 | 0.08 |
| | | | Partis | 68327 | 10095 | 0.95 | 0.88 | 0.92 |
| | | | SCOPe | 67312 | 7192 | 0.96 | 0.89 | 0.93 |

Table 17: Comparison of MobiLLE with four different clonal grouping methods on the three experimental data sets, using **closeness evaluation method.**

| | # seq | # clones by MobiLLe | | # analyzed seq | # clones | Precision | Recall | F-score |
|---|---|---|---|---|---|---|---|---|
| $I_1$ | 33599 | 162 | BRILIA | 33578 | 161 | 1 | 0.06 | 0.11 |
| | | | SONAR | 29335 | 3542 | 0.99 | 0 | 0 |
| | | | Partis | 33585 | 73 | 1 | 0.14 | 0.25 |
| | | | SCOPe | 33554 | 176 | 1 | 0.01 | 0.02 |
| $I_2$ | 140026 | 2398 | BRILIA | 90236 | 7234 | 0.99 | 0.01 | 0.01 |
| | | | SONAR | 138439 | 1975 | 0.99 | 0.19 | 0.32 |
| | | | Partis | 138688 | 927 | 0.98 | 0.66 | 0.79 |
| | | | SCOPe | 138250 | 1387 | 0.97 | 0.5 | 0.66 |
| $I_8$ | 70050 | 11256 | BRILIA | 68133 | 10461 | 0.96 | 0.76 | 0.85 |
| | | | SONAR | 68171 | 17096 | 0.43 | 0.07 | 0.12 |
| | | | Partis | 68327 | 10095 | 0.94 | 0.68 | 0.79 |
| | | | SCOPe | 67312 | 7192 | 0.89 | 0.77 | 0.83 |

Table 18: Comparison of MobiLLE computational time (in seconds) with four different clonal grouping methods on the three experimental data sets.

| | MobiLLe | Partis | SONAR | BRILIA | SCOPe |
|---|---|---|---|---|---|
| $I_1$ | 17 | 2580 | 19 | 253 | 109 |
| $I_2$ | 208 | 5104 | 265 | 552 | 330 |
| $I_8$ | 135 | 1562 | 187 | 446 | 187 |

#### 1.1 Figures

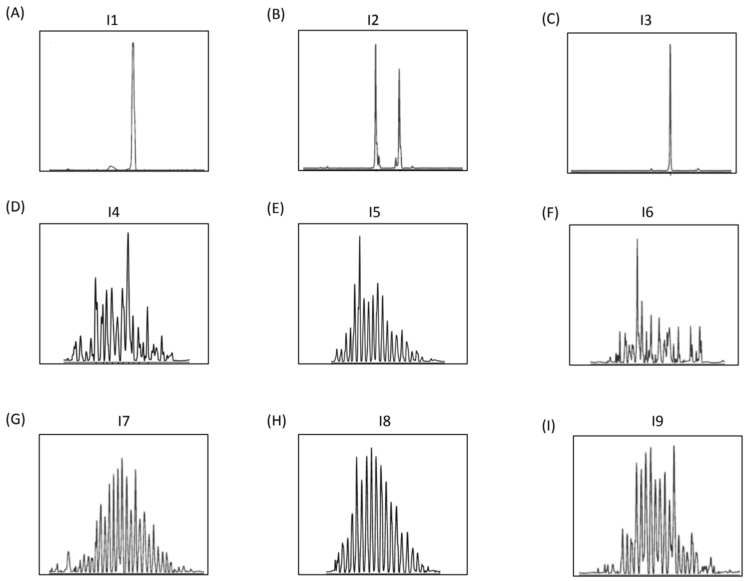


Figure 1: **GeneScan profiles of human peripheral blood samples**. IGH-VDJ rearrangements were amplified using conventional methods and PCR products were further analyzed by capillary electrophoresis. (A-C) Samples from individuals with monoclonal B-cell malignancy: monoallelic profile (A and C) or biallelic profile (B); (D-I) non-malignant samples: regular polyclonal profile (D, E, G, H, I) or irregular polyclonal profile (F).

**Legend for Figures S3 to S11**

A) Circle representation of the clone abundance. Each circle symbolizes a clone, and the clone's abundance is shown through its size.

B) Number of sequences in each clone, all clones are represented, vertical axe is in log scale.

C) Lorenz curve and Gini coefficient. A Lorenz curve shows the graphical representation of clonal inequality. On the horizontal axe, it plots the cumulative fraction of total clones when ordered from the less to the most abundant; On the vertical axe,it show the cumulative fraction of sequences.

D) Clonal size distribution (percentage) of the 100 most abundant clones.

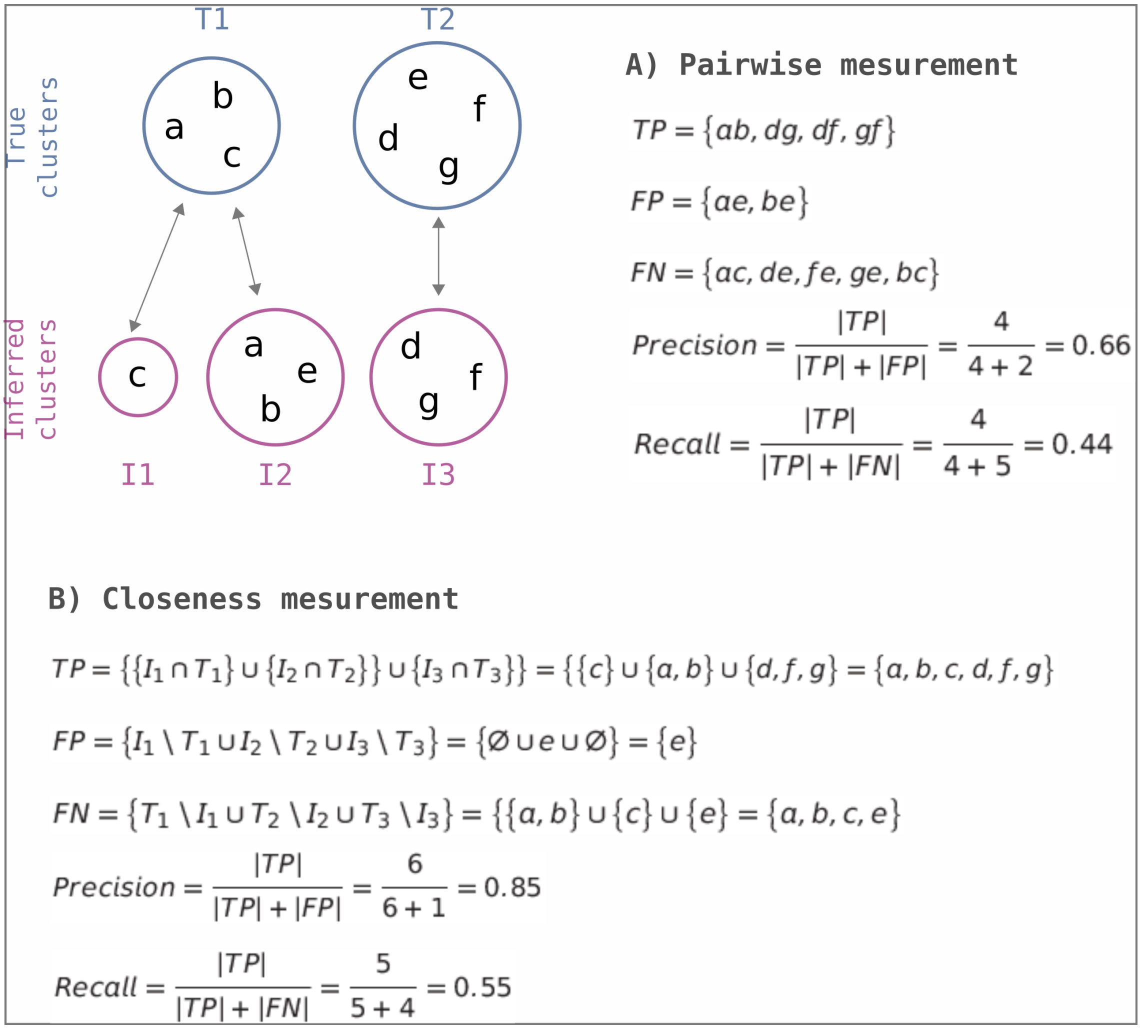


Figure 2: **Clustering performance measurements.** A) Pairwise B) Closeness.

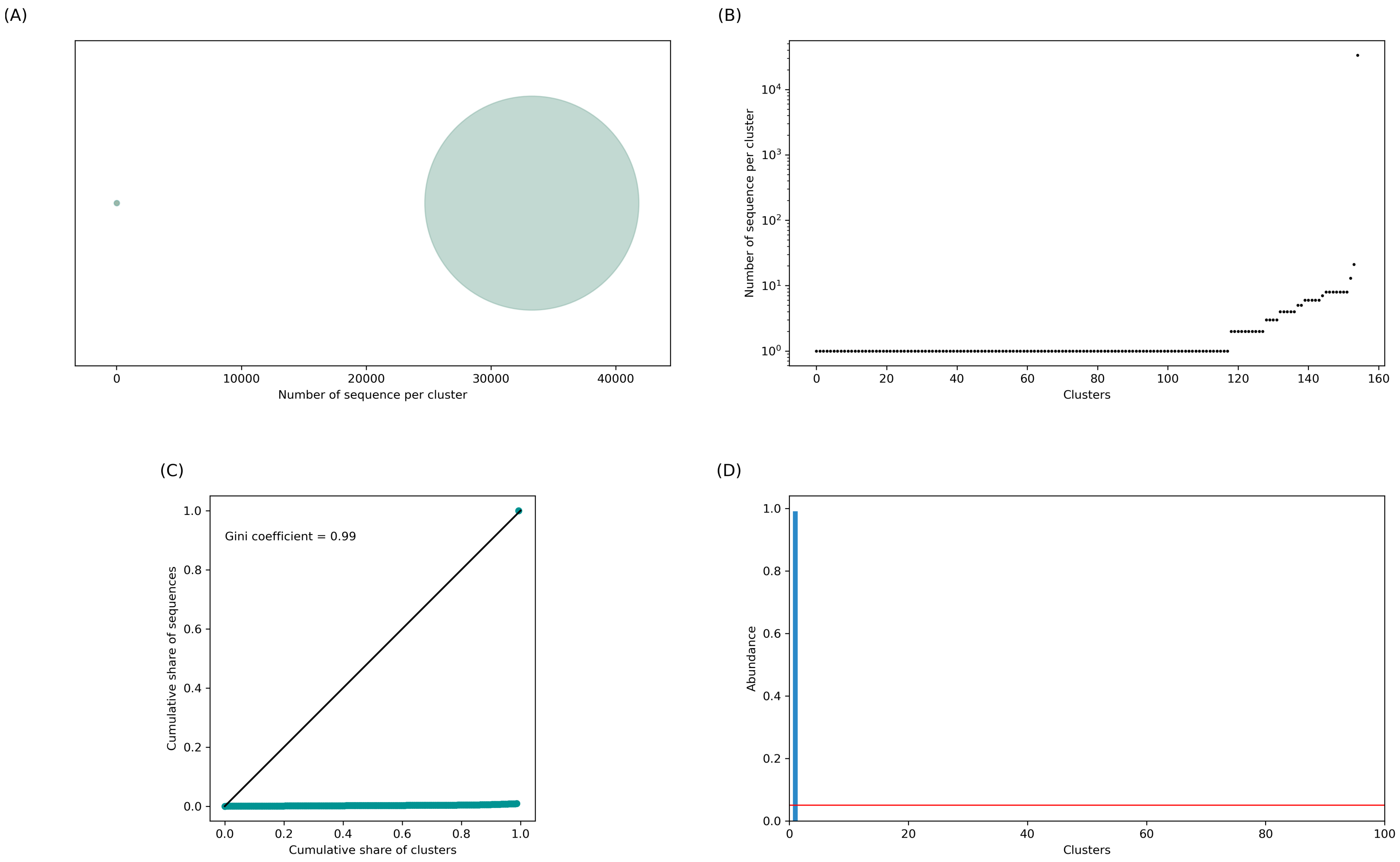


Figure 3: **Repertoire of individual 1**

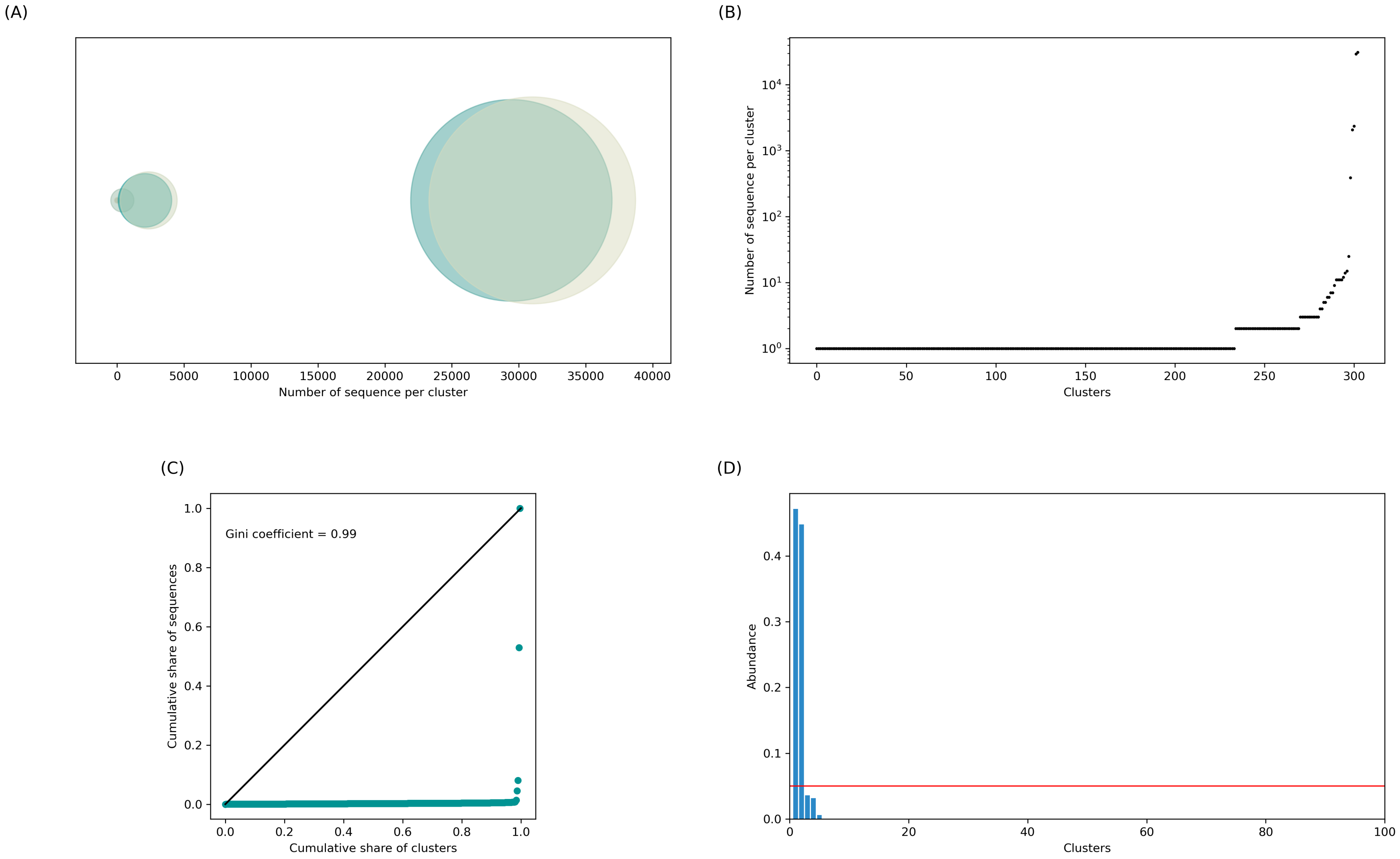


Figure 4: **Repertoire of individual 2.**

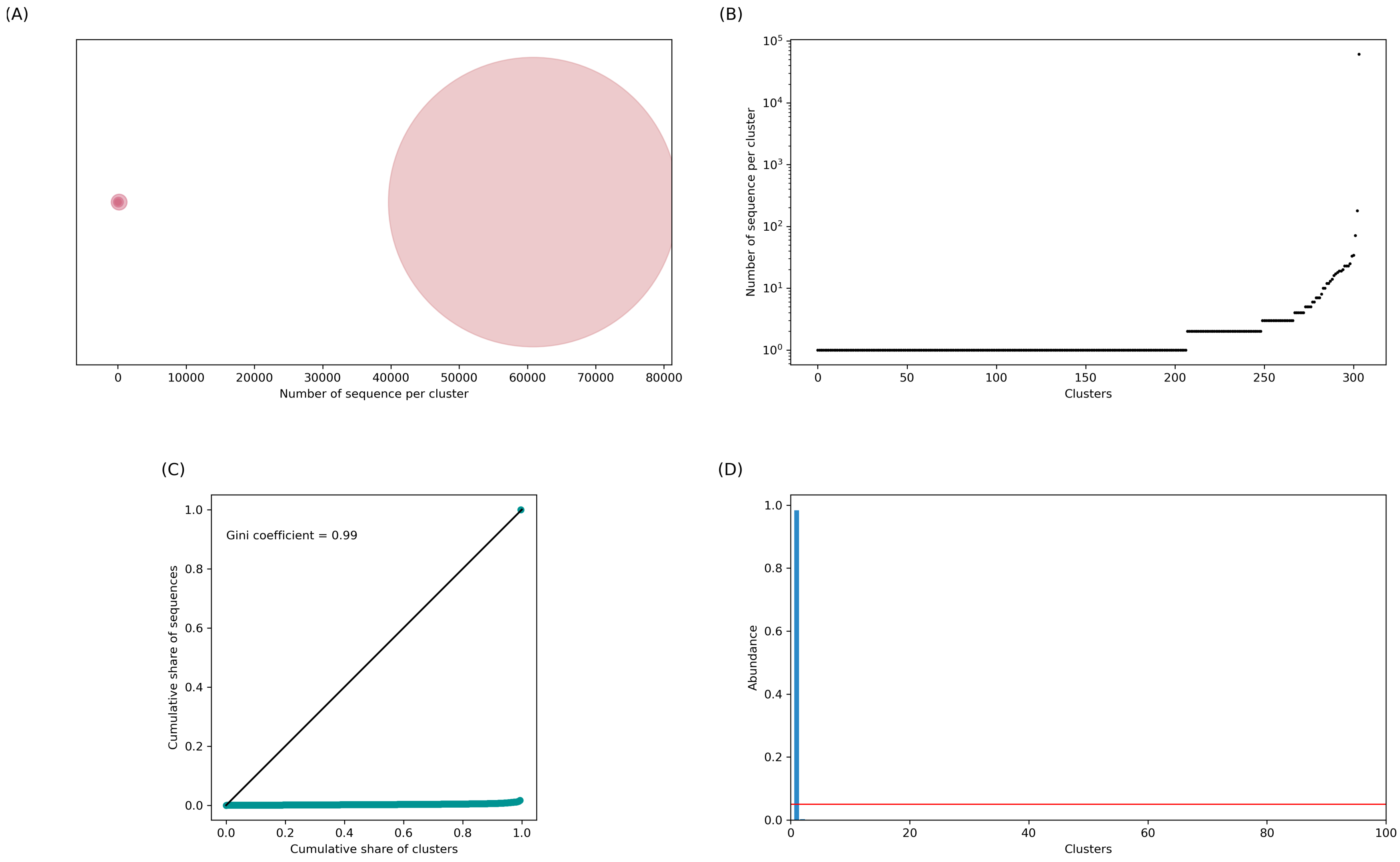


Figure 5: **Repertoire of individual 3**.

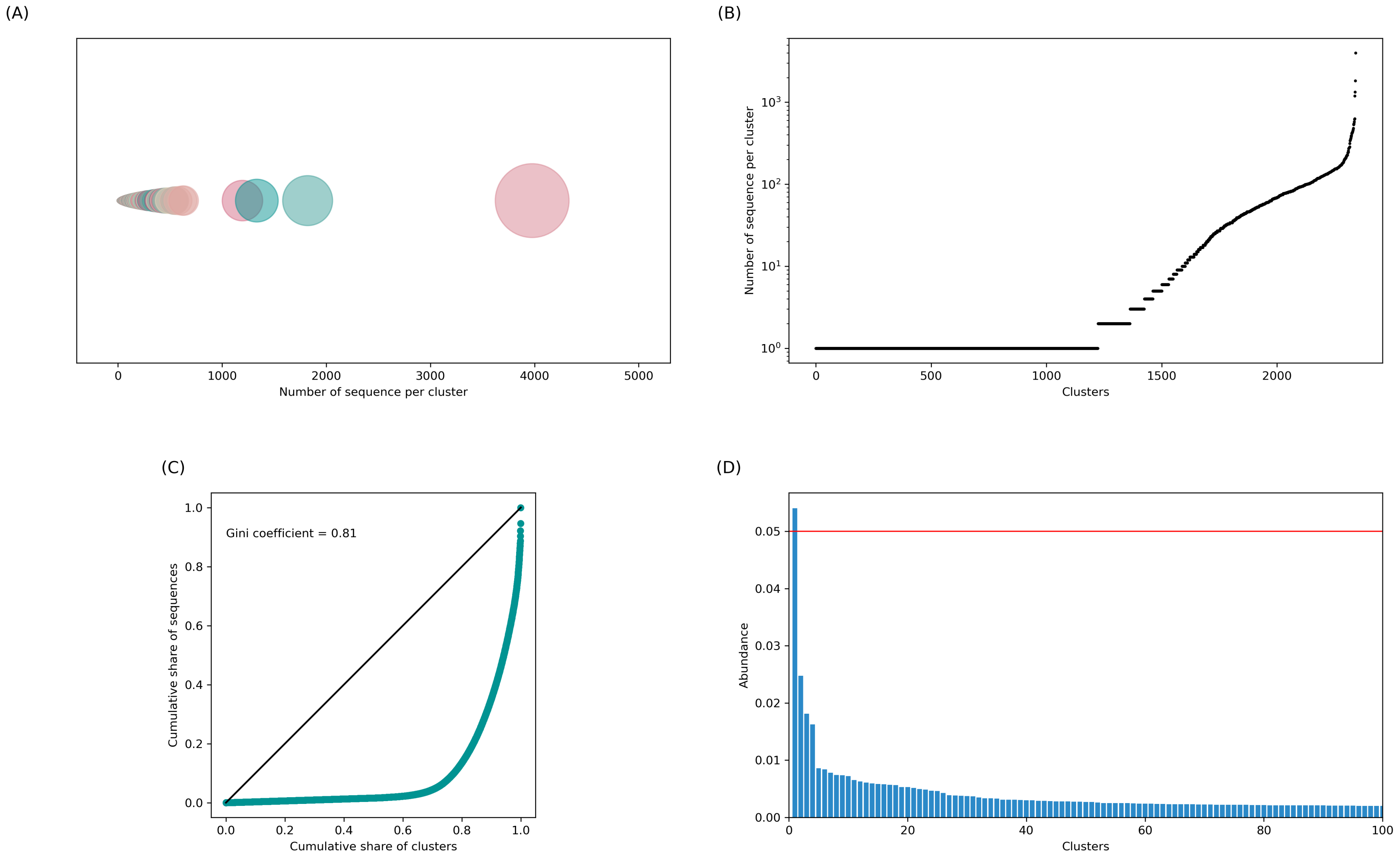


Figure 6: **Repertoire of individual 4.**

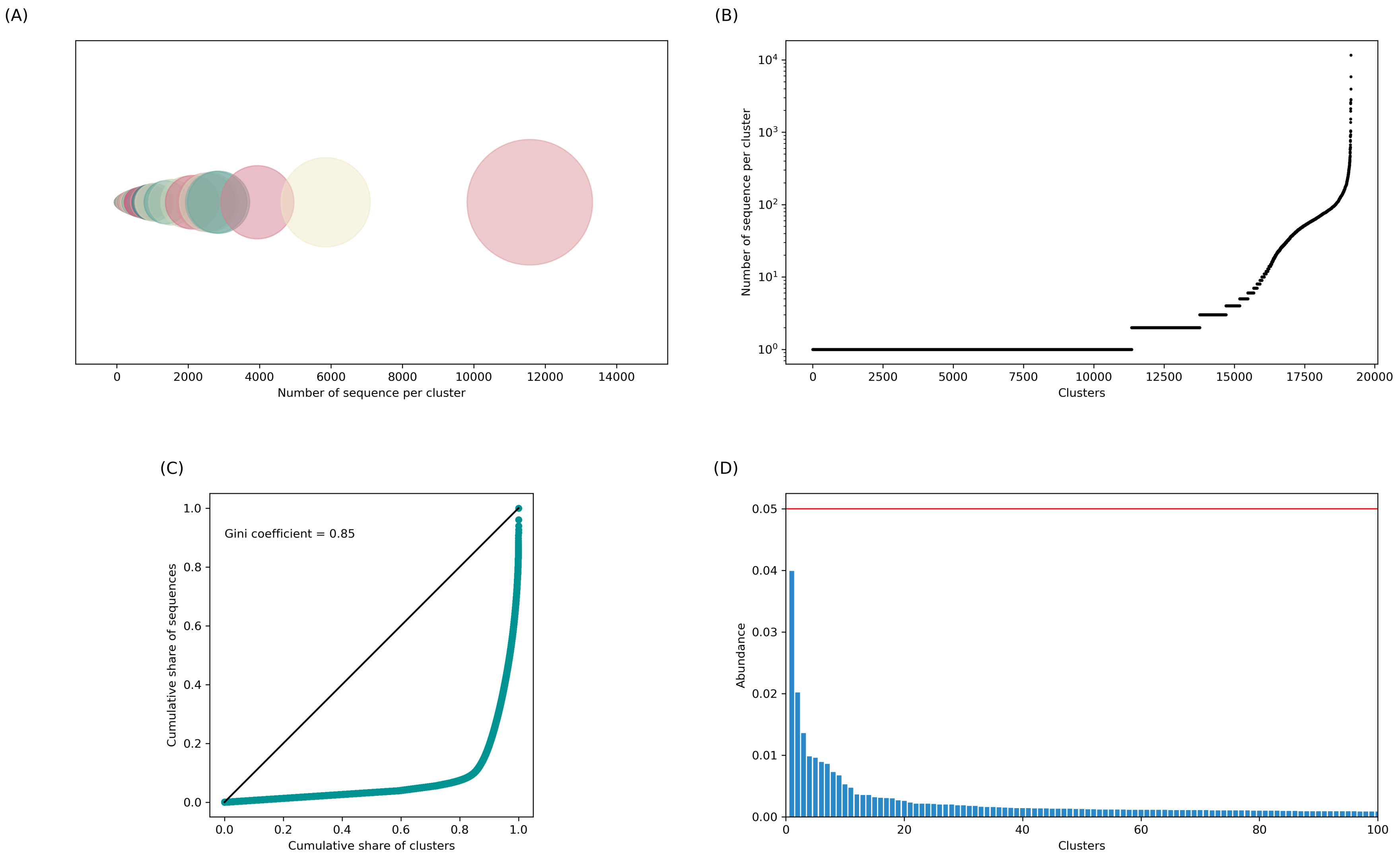


Figure 7: **Repertoire of individual 5.**

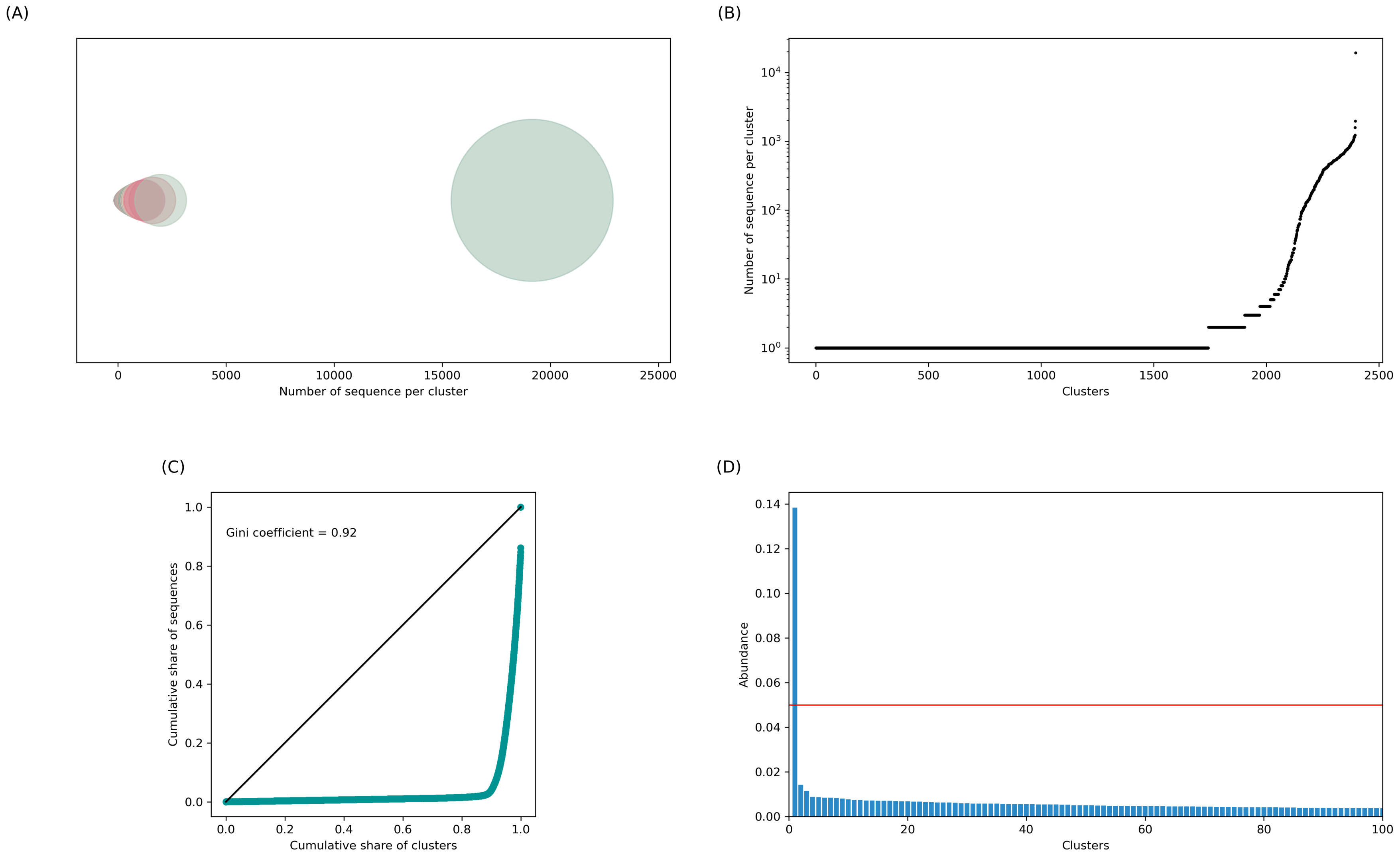


Figure 8: **Repertoire of individual 6.**

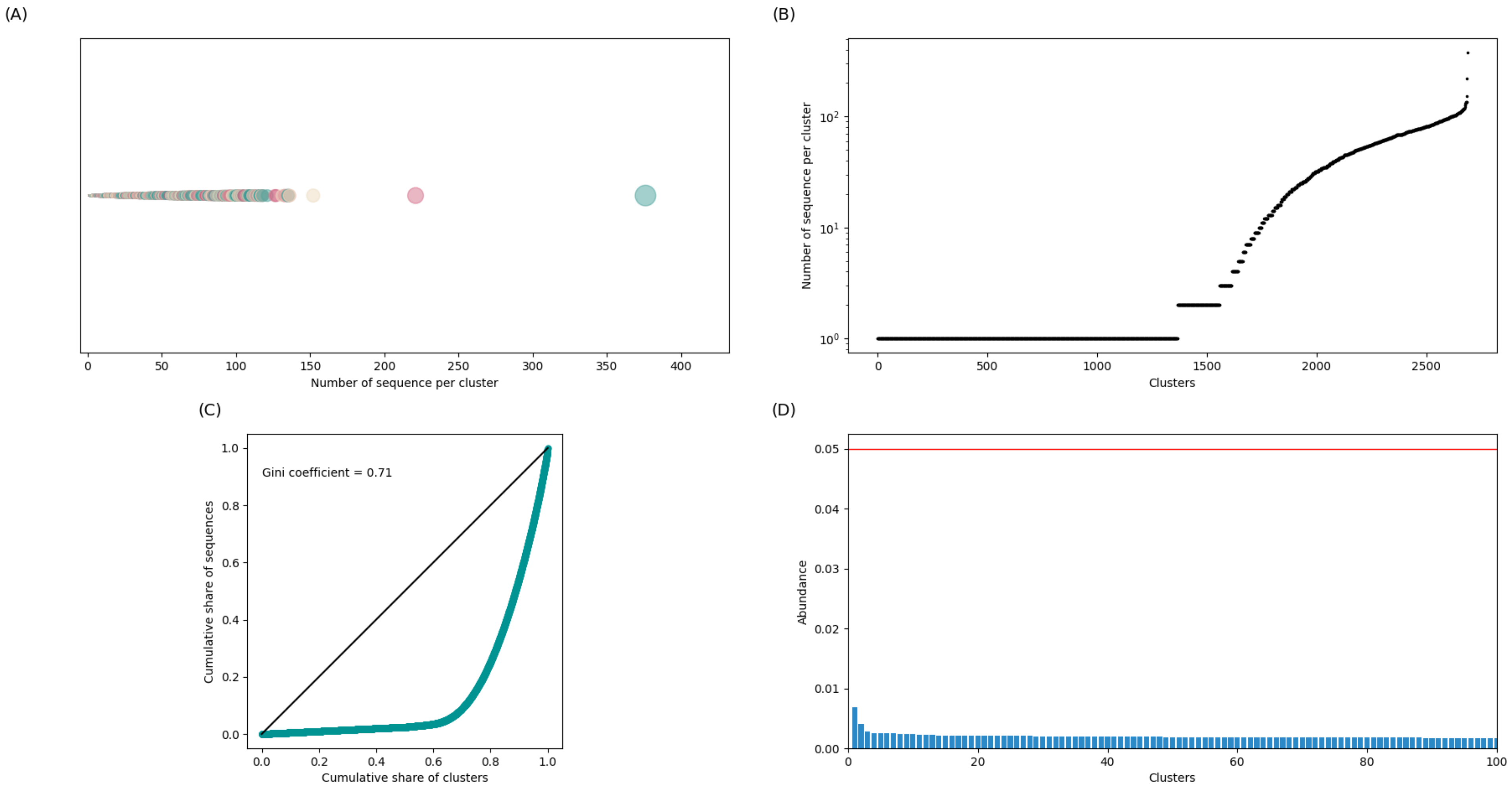


Figure 9: **Repertoire of individual 7**.

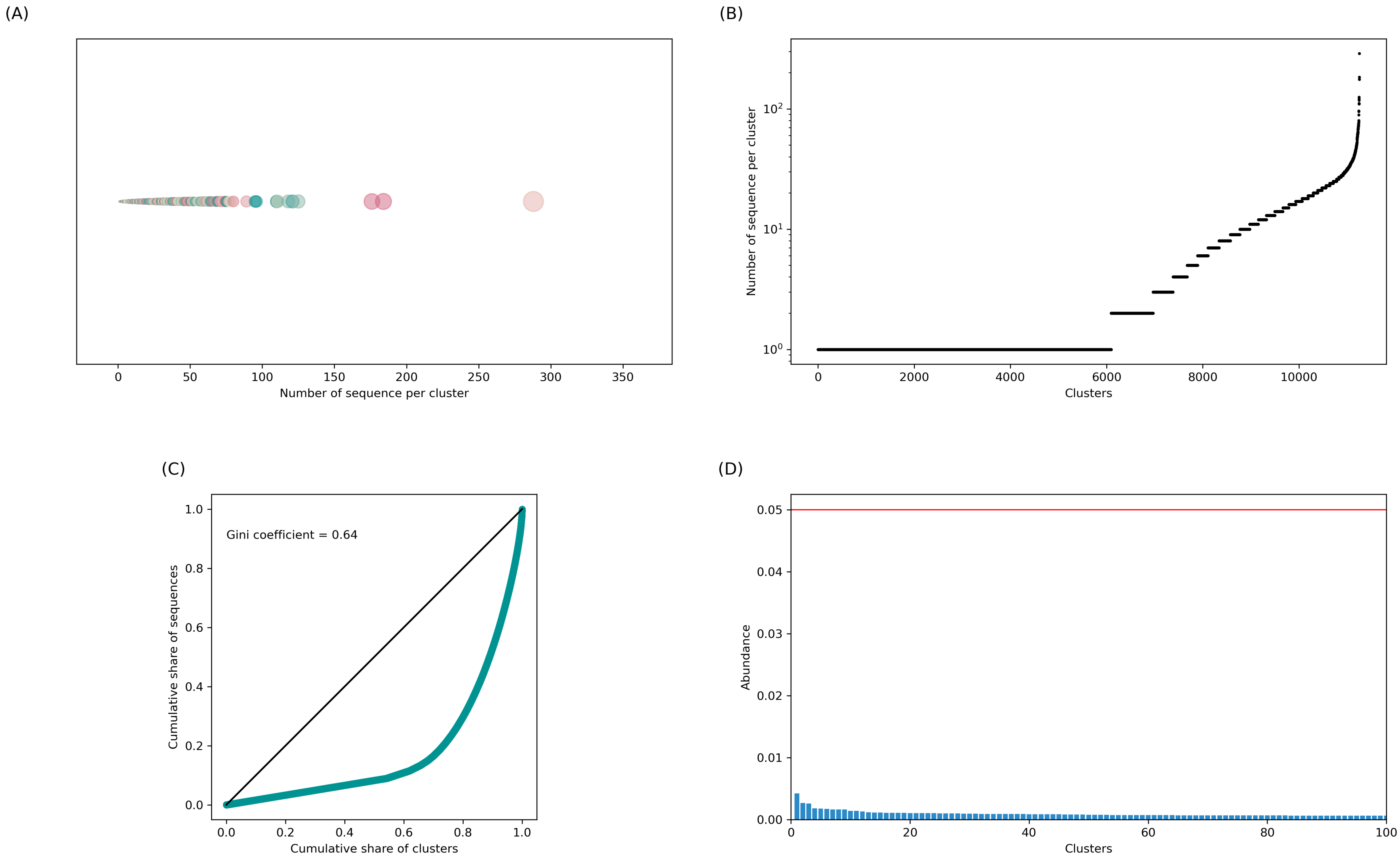


Figure 10: **Repertoire of individual 8.**

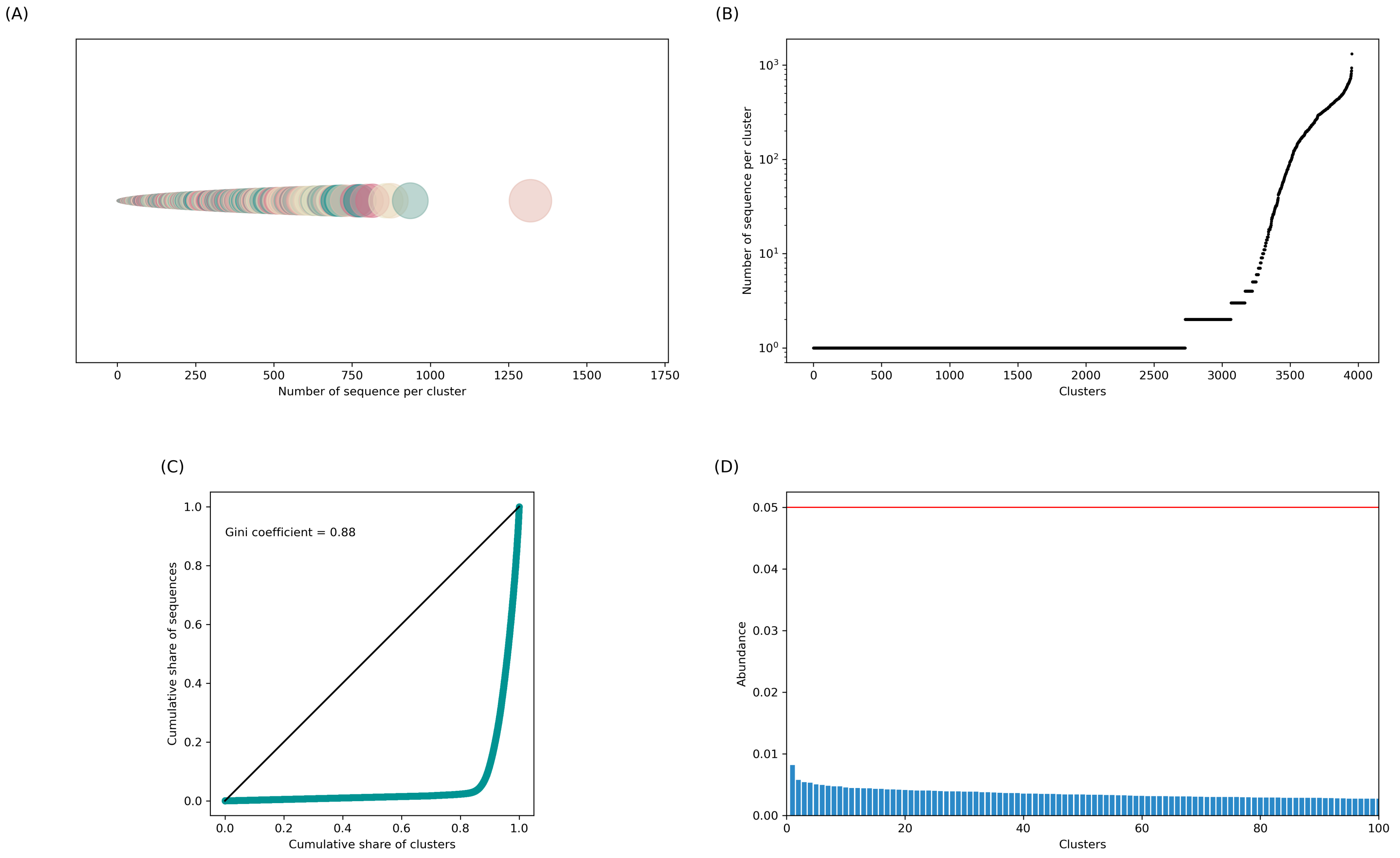


Figure 11: **Repertoire of individual 9.**
